# Fetal sex shapes placental inflammatory responses to extracellular mitochondrial DNA

**DOI:** 10.64898/2026.07.09.737607

**Authors:** Reneé de Nazaré Oliveira da Silva, Nataliia Hula, Desirae Escalera, Leslie Lopez, Gabrielle Kelly, Isabelle K. Gorham, Megan Rowe, Contessa A. Ricci, Ciprian Gheorghe, Nicole R. Phillips, Styliani Goulopoulou

## Abstract

Aberrant changes in circulating cell-free mitochondrial DNA (ccf-mtDNA) across gestation are associated with adverse pregnancy outcomes. Given the inflammatory properties of ccf-mtDNA via pattern recognition receptors such as Toll-like receptor 9 (TLR9), we hypothesized that extracellular mtDNA induces placental inflammation via TLR9 signaling and that this response differs by fetal sex. Pregnant Sprague-Dawley rats were treated intravenously with purified mtDNA (300 μg/kg), nuclear DNA (nDNA), saline, and/or the TLR9 antagonist ODN2088 across five studies. Placental responses were evaluated 4 h (Studies 1-3) and 24 h (Study 4) post-treatment; pregnancy and neonatal outcomes were assessed at delivery (Study 5). Exposure to mtDNA, but not nDNA, increased placental *il1β*, *tnf⍺*, and *il10* mRNA (p < 0.05), establishing response specificity. mtDNA-induced placental inflammation was fetal sex-dependent: mtDNA increased *il6* and *il1β* mRNA in male placentas (p ≤ 0.0004) but not female placentas, whereas *ifnγ* was selectively induced in female placentas (p = 0.0004). TLR9 and MyD88 abundance increased in female but not male placentas, and TLR9 antagonism modified selected inflammatory responses with sex-specific patterns. The 4 h inflammatory transcriptional signature resolved by 24 h, whereas mtDNA exposure was associated with a sex-specific shift in antioxidant enzyme expression persisting to 24 h. Despite no effects on gestational length or neonatal biometrics, mtDNA exposure was associated with a higher estimated stillbirth count per litter (IRR = 4.23, 95% CI [0.89, 20.1], p = 0.069). These findings establish extracellular mtDNA as an acute, sex-differentiated placental inflammatory stimulus with partial TLR9 dependence and a potential impact on fetal viability.

**New & Noteworthy:** This study demonstrates that acute exposure to extracellular mtDNA induces placental inflammatory responses in vivo. This response is specific to mtDNA, fetal-sex dependent, and partially mediated by TLR9, with male and female placentas engaging distinct inflammatory signals within hours of exposure. The biological effects extend beyond the initial inflammatory window, with mtDNA exposure producing lasting, sex-specific changes in antioxidant enzyme expression. mtDNA-exposed dams had higher expected stillbirth counts, suggesting extracellular mtDNA may affect fetal viability.

## Introduction

The placenta is the critical interface between mother and fetus, supporting fetal growth by supplying oxygen and nutrients while coordinating immune and endocrine signaling (1, 2). In mammals, its hemochorial architecture, in which the chorion contacts maternal blood directly, provides a thin, specialized barrier enabling efficient maternal-fetal exchange while regulating transport (3). However, during pregnancies complicated by inflammation, such as preeclampsia, the placenta releases proinflammatory, vasoactive, and antiangiogenic factors, and due to its hemochorial structure, it facilitates their rapid access to maternal blood (4). Once in the maternal bloodstream, these factors interact with maternal organs, amplifying widespread systemic inflammation.

Circulating cell-free mitochondrial DNA (ccf-mtDNA) is a damage-associated molecular pattern (DAMP) recognized by innate immune pattern-recognition receptors (5). In healthy pregnancy, ccf-mtDNA is detectable in maternal blood, increases with advancing gestational age, and returns to pre-pregnancy levels after delivery (6). The normal gestational trajectory of ccf-mtDNA is disrupted in adverse pregnancy outcomes associated with inflammation(7–10). Given that ccf-mtDNA is recognized by innate immune pattern recognition receptors, dysregulation of its gestational profile has the potential to contribute to sterile inflammatory signaling, defined as innate immune activation in the absence of an infectious trigger.

Extracellular mtDNA is recognized by Toll-like receptor 9 (TLR9), triggering intracellular signaling cascades that promote nuclear factor kappa B (NF-κB) nuclear translocation and transcription of proinflammatory mediators, including monocyte chemoattractant protein-1 (MCP-1), interleukins (IL-1β and IL-6), granulocyte–macrophage colony-stimulating factor (GM-CSF), and tumor necrosis factor-alpha (TNF-α) (5, 11). In humans, placental TLR9 expression and signaling activity are increased in pregnancies with preeclampsia, a hypertensive disorder of pregnancy associated with systemic inflammation (12–14). In rodent models, systemic TLR9 stimulation during pregnancy induces placental inflammation and causes adverse fetoplacental outcomes (15, 16). Nevertheless, whether and how recognition of ccf-mtDNA by TLR9 drives placental inflammation during pregnancy, particularly in a fetal sex-dependent manner, remains poorly defined.

Fetal sex shapes placental responses to inflammatory and stress-related stimuli and is associated with pregnancy complications including gestational diabetes, preterm birth (17–19), placental abruption (20), and stillbirth (21). Consistent with sex-dependent immune regulation, maternal psychosocial stress is associated with higher circulating IL-1β and IL-6 in women carrying male fetuses compared to female fetuses (22), and exposure to lipopolysaccharide (LPS) increases TLR expression in trophoblast cells derived from male placentas (23).

These sex-dependent differences in placental immune responsiveness may reflect multiple, non-mutually exclusive mechanisms. First, sex chromosome complement influences immune gene dosage. Several immune-relevant genes located on the X chromosome escape X-inactivation, leading to higher expression in XX than in XY cells. This increased gene dosage may influence innate immune signal transduction in female placentas (24, 25). Second, gonadal hormones differentially modulate TLR-mediated innate immune responses. Estrogen enhances TLR9-mediated signaling and downstream NF-κB activation in immune cells (26), whereas androgens suppress TLR-mediated proinflammatory responses in macrophages and other immune cell types (27). Third, baseline differences in placental immune cell composition and inflammatory tone between male and female placentas may establish differing activation thresholds that determine the magnitude and profile of responses to innate immune triggers (22). Together, these mechanisms provide a biological rationale for expecting that the placental response to extracellular mtDNA would differ by fetal sex. Therefore, we hypothesized that exposure to extracellular mtDNA during pregnancy induces placental inflammation via TLR9 signaling and that the magnitude and profile of this response differ by fetal sex.

## Methods

### Chemicals and Reagents

Details of chemicals and reagents used in this study are provided in Supplementary File 1 **(Supplementary File 1: Table S1)**.

### Animals

All animal protocols and procedures were approved by the Institutional Animal Care and Use Committees (IACUC) of Loma Linda University (IACUC-22-003) and were performed in accordance with the Guide for the Care and Use of Laboratory Animals of the National Institutes of Health.

Sprague-Dawley rats were purchased from Envigo (Indianapolis, IN, USA). Male rats (body weight and age on arrival: ∼400 g and 13-15-weeks old) were used for breeding purposes. Female rats were purchased either timed-pregnant and arrived at the animal facilities on gestational day (GD) 5-8 (body weight and age on arrival: ∼220 g and 11-12-week-old), with gestational age determined by Envigo based on the presence of a vaginal plug or were bred in-house as previously described (15, 28). Female rats used for breeding weighed 190-230 g and were 9-11-week-old at arrival. GD1 was designated as the day on which spermatozoa were observed in vaginal smears after pair mating.

Rats were pair-housed under 12:12-h light/dark cycles (lights on, 07:00 h; lights off, 19:00 h) in a temperature– and humidity-controlled environment. Animals were allowed to acclimatize for one week before handling and were provided standard laboratory chow and water ad libitum throughout the study. All experiments were performed when rats were 12-24 weeks old. In total, 140 female and 8 male rats were used. Sample sizes were selected based on our previous studies using similar animal models, interventions, and outcome measures (15, 29).

### Experimental design, timeline, and treatments

This study is part of a larger research project evaluating the effects of extracellular mtDNA on maternal vascular and fetoplacental responses. Placental outcomes reported here were derived from two sources: a) cohorts that overlap with those from which maternal vascular tissues were collected for vascular function assessments and reported separately (30). Specifically, the same animals contributed both placental tissue for the current study and maternal vascular tissues for vascular function analyses reported elsewhere (see Studies 1-3 below); and b) additional, non-overlapping cohorts were collected under the same experimental framework to capture placental endpoints and pregnancy outcomes requiring distinct collection conditions (see Studies 4-5 below). Separate cohorts of animals were used in each study described in this manuscript. Treatments were consistent across all study arms; however, not all animals contributed data to both manuscripts.

Five studies were performed, each with a separate cohort of animals **(Figure 1)**: In *Study 1*, we established the specificity of mtDNA in inducing pro-inflammatory responses in placenta. Pregnant rats were treated with purified mtDNA (300 µg/kg body weight; mtDNA group), sterile 0.9% saline (vehicle; Saline group), or nuclear DNA (300 µg/kg body weight; nDNA group).

**Figure 1.**
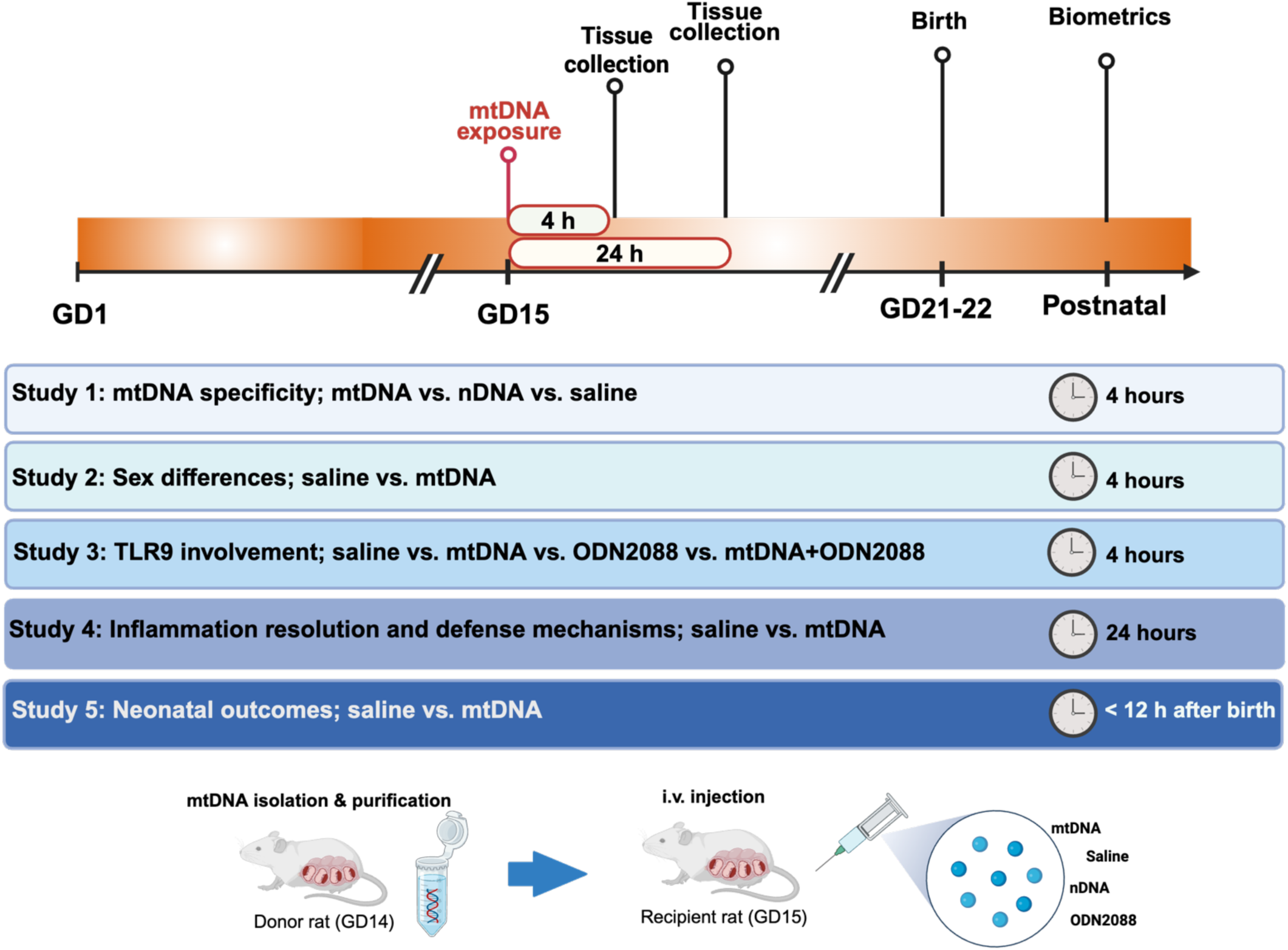
Experimental design. Experimental design of studies evaluating the effects of extracellular mitochondrial DNA (mtDNA) exposure on placental inflammatory responses, inflammation resolution, TLR9 signaling, pregnancy outcomes, and neonatal outcomes in pregnant rats. Pregnant rats were treated intravenously with purified mtDNA, nuclear DNA (nDNA), saline vehicle, and/or the TLR9 antagonist ODN2088 depending on study design. Studies 1-3 evaluated acute responses 4 h after treatment, Study 4 evaluated responses 24 h after treatment, and Study 5 assessed pregnancy and neonatal outcomes at delivery. Separate cohorts of animals were used for each study. Created with Biorender.com.

In *Study 2*, we determined the impact of fetal sex on placental inflammatory responses to acute exposure (4 h) to purified mtDNA. Pregnant rats were treated with mtDNA (300 µg/kg body weight; mtDNA group) or sterile 0.9% saline (vehicle; Saline group).

In *Study 3*, we investigated whether mtDNA-induced placental inflammatory responses were mediated via TLR9 signaling. Pregnant rats were treated with mtDNA (300 µg/kg body weight; mtDNA group), sterile 0.9% saline (vehicle; Saline group), ODN 2088 (60 µg/kg body weight) plus sterile 0.9% saline (ODN2088 group), or ODN 2088 (60 µg/kg body weight) plus mtDNA (300 µg/kg body weight; ODN2088+mtDNA group). ODN2088 (synthetic TLR9-specific antagonist) was administered via an i.v. injection (tail vein) 5 min before saline and mtDNA injections.

In *Study 4*, we examined whether the placental inflammatory responses to purified mtDNA were resolved 24 h post-treatment and if inflammation resolution differ between placentas from male and female fetuses. Pregnant rats were treated with mtDNA (300 µg/kg body weight; mtDNA group) or sterile 0.9% saline (vehicle; Saline group).

In *Study 5,* we assessed the effects of gestational exposure to purified mtDNA on pregnancy and neonatal outcomes. Pregnant rats were treated with mtDNA (300 µg/kg body weight; mtDNA group) or sterile 0.9% saline (vehicle; Saline group). After treatments, rats were returned to clean cages for standard post-anesthesia monitoring and then moved back to animal facilities until delivery. Pregnancy outcomes (litter size: live births, stillborn pups; gestational length) and neonatal biometrics (neonatal weight, crown-to-rump length, and abdominal girth) were recorded within 12 h after delivery.

Study 1 included one randomly selected placenta per litter. In Studies 2-4, one placenta associated with a male fetus and one placenta associated with a female fetus were collected from each litter for analysis.

In all studies, pharmacological treatments were delivered via intravenous (i.v.; tail vein) injections, which were administered within 1 min using a 1 mL syringe with 25G x 5/8’’ needle while rats were under isoflurane anesthesia (5% for induction, 3% for maintenance, 100% oxygen). Treatments were delivered at 8:00 – 9:00 h.

After treatments, rats were placed in clean cages and monitored until euthanasia (Studies 1-3) or until full recovery (Studies 4-5). Rats were euthanized 4 h after treatment (Study 1-3) or 24 h after treatment (Study 4). Animals in Study 5 were not euthanized at a fixed timepoint but were allowed to deliver at term, at which time neonatal biometrics were recorded.

Purified mtDNA and nuclear DNA (nDNA) were isolated from liver of pregnant donor rats (GD14) as described in our companion vascular outcomes manuscript (30). The liver was selected because of its high mitochondrial content and prior use in rodent models of mitochondrial DAMP-induced inflammation (5). The mtDNA dose was based on pilot studies and previously published work demonstrating the biological activity of exogenous mtDNA in rodent models (5). Working solutions of ODN2088 were prepared in endotoxin-free water, followed by dilution in sterile 0.9% saline immediately before use.

Experiments were conducted in pregnant rats at GD14-15. This gestational window was selected because earlier immune stimulation can disrupt placental development and pregnancy viability (16, 31), while GD14-15 represents a period of rapid fetoplacental growth (32, 33) and is consistent with our previous studies of innate immune system activation during pregnancy (15).

### Euthanasia and tissue collection

Rats were anesthetized with isoflurane (5% for induction, 3% for maintenance, 100% oxygen) and euthanized by isoflurane overdose, followed by bilateral thoracotomy and removal of the heart. Prior to euthanasia and when rats were under a deep plane of anesthesia, whole blood was collected from the inferior vena cava into EDTA-coated collection tubes (BD, Franklin Lakes, NJ; Cat No. 367856). Plasma was isolated from whole blood by centrifugation (2,000 × g, 15 min, 4°C), snap-frozen in liquid nitrogen, and stored at –80°C for subsequent quantification of ccf-mtDNA and measurements of circulating cytokines. After euthanasia, corresponding fetal and placental samples from both left and right uterine horns were excised, washed with phosphate-buffered saline, and immediately snap-frozen and stored at –80°C for fetal-placental sex determination assays and molecular assessment of protein and RNA levels with Western blotting and qPCR, respectively.

### Donor tissue, mitochondrial and nuclear enrichment, and DNA extraction

Detailed description of isolation of purified mtDNA and nDNA from rat liver tissues is published in our companion vascular outcomes manuscript (30). Briefly, mitochondria were enriched from 100-200 mg liver tissue using a commercially available mitochondria isolation kit (Thermo Scientific). The obtained mitochondrial pellet was then used for mtDNA extraction. In parallel, nuclei were enriched from 1 g liver by sucrose-based fractionation. The resulting pellet was then subjected to a DNA extraction protocol. DNA was isolated from mitochondrial and nuclear pellets using the QIAamp DNA Mini Kit (QIAGEN LLC) according to the manufacturer’s instructions. DNA concentrations were quantified by spectrophotometry (NanoDrop™ OneC, Thermo Fisher Scientific, Waltham, MA, USA), and DNA was stored at 4 °C and used within 1 month of extraction.

Quality control data from mtDNA purity and integrity assessments have been included in our published companion vascular outcomes manuscript (30). Briefly, mtDNA purity and integrity were confirmed by spectrophotometry (A260/280), bicinchoninic acid (BCA) protein assay, qPCR-based assessment of mtDNA enrichment relative to nDNA (to estimate nDNA carryover), chromogenic Limulus amebocyte lysate (LAL) assay (Thermo Fisher Scientific), and fragment sizing by TapeStation.

### Circulating cell-free mitochondrial DNA quantification

DNA was extracted from 200 µL maternal plasma using the DNeasy Blood and Tissue Kit (QIAGEN) according to the manufacturer’s instructions and our previous publications (29), starting at the lysis buffer step. DNA was eluted in 200 µL of DNase-free water and concentrated from a volume of 200 µL to a volume of 20 µL using the Vacufuge vacuum concentrator (Eppendorf).

Nuclear DNA and mtDNA were quantified by quantitative PCR (qPCR) on a 7500 Real-Time PCR System (Applied Biosystems™, Waltham, MA, USA), as we previously described (29). TaqMan assays targeting the mitochondrial D-loop and the β-actin gene were used to measured mtDNA and nDNA, respectively. For each reaction, the qPCR master mix contained 13 µL of TaqMan^TM^ Universal Master Mix II, no UNG (Applied Biosystems™), 2 µL of each primer (D-loop, 0.625 µM; β-actin, 2.5 µM) and 1 µL of probe (D-loop, 2.5 µM; β-actin, 2.5 µM). Reactions were assembled in a 96-well plate with 2 µL of DNA sample or DNase-free water to serve as a negative control.

mtDNA copy number was determined using a standard curve generated from a synthetic mtDNA fragment of known copy numbers (Sequence: GGTTCTTACTTCAGGGCCATCAATTGGTTCATCGTCCATACGTTCCCCTTAAATAAGACATCTCGATGGTAACGGGTCTAATC) included on every plate. Standard curve linearity (R^2^) and amplification efficiency were assessed for each run to ensure assay performance. Negative controls were included in every run to monitor nonspecific mtDNA amplification and confirm assay specificity. mtDNA concentrations are reported as copies/mL plasma. The targeted sequences for primers and probes were:

- Mitochondrial D-loop. Forward: GGTTCTTACTTCAGGGCCATCA; Reverse: GATTAGACCCGTTACCATCGAGAT; Probe: 6FAM-TTGGTTCATCGTCCATACGTTCCCCTTA-TAMRA, (GenBank accession no. X14848)
- Nuclear target β-actin. Forward: GGGATGTTTGCTCCAACCAA; Reverse: GCGCTTTTGACTCAAGGATTTAA; Probe: VIC-CGGTCGCCTTCACCGTTCCAGTT-TAMRA, (GenBank accession no. V01217)

### Plasma and placental cytokine measurements

Plasma and placental cytokines/chemokines were measured using the Bio-Plex™ Rat Cytokine/Chemokine Magnetic Bead Panel (Cat. #64511854, Bio-Rad, Foster City, CA, USA) on a Bio-Plex® 200 System MRW A534 (Bio-Rad, Foster City, CA, USA), following the manufacturer’s instructions. Samples, standards, and quality controls were run in duplicate. Plasma was diluted 1:4 and placental lysates were diluted 1:2 in assay buffer. Total protein concentrations in placental lysates were determined using the Pierce™ BCA Protein Assay. Only analytes with concentrations falling within the manufacturer-specified assay ranges were included in the analysis and are reported herein.

### Placental sex determination

Placental sex was determined using a single-step PCR assay as previously described by Dhakal et al. (34) and adapted in our prior work (29). Briefly, DNA was isolated from fetal tissue and amplified using three primers targeting sex-specific sequences **(Supplementary File 1: Table S2)**. PCR products were separated on a 1% agarose gel containing ethidium bromide. PCR products from tail DNA of adult male and female rats served as positive controls. Gels were imaged under ultraviolet illumination using Azure300 imaging system (Azure Biosystems, Dublin, CA). Images from sex determination assays are provided in **Supplementary File 2**.

### RNA isolation, cDNA synthesis, and quantitative real-time polymerase chain reaction

Total RNA was extracted from placental tissue (100 mg) using QIAzol® Lysis Reagent (QIAGEN) and the miRNeasy Mini Kit (QIAGEN) according to the manufacturer instructions. RNA concentration and purity were assessed by spectrophotometry (NanoDrop^TM^ OneC), cDNA was synthesized using Sensiscript RT Kit (QIAGEN) supplemented with RiboGuard RNase inhibitor (LGC, Biosearch Technologies) and oligo(dT) primers (QIAGEN), following our published protocols (15, 29).

Placental mRNA expression of pro-inflammatory and anti-inflammatory cytokines, immune cell markers, and *gapdh* (reference gene) was quantified by quantitative real-time polymerase chain reaction (qRT-PCR) using SYBR Green chemistry on a CFX96 Real-Time PCR Detection System (Bio-Rad). Primer sequences are provided in **Supplementary File 1 (Table S3)**. All targets were analyzed in duplicate and melt curve analysis was performed to confirm amplification specificity. Relative gene expression was calculated using the 2^-ΔΔCT^ method with *gapdh* used for normalization.

### Western blot analysis

Protein lysates were prepared from placental tissue via T-PER^TM^ Tissue Protein Extraction Reagent (Thermo Fisher Scientific) supplemented with protease inhibitors (Sigma Aldrich). Total protein concentrations were measured with the Pierce^TM^ BCA Protein assay. Equal protein amounts (15-30 µg) were separated by SDS-PAGE and transferred to nitrocellulose membranes (Bio-Rad) using a Trans-Blot^®^ Turbo^TM^ system (Bio-Rad) for 10 min. Membranes were blocked (1 h) in either 3% bovine serum albumin (BSA; Sigma-Aldrich) or 5% non-fat dry milk (Bio-Rad) prepared in tris-buffered saline with Tween-20 (T-BST), then incubated overnight at 4°C with primary antibodies. Protein signals were detected using Odyssey CLx (LI-COR Biosciences, Lincoln, NE, USA) or AZURE 300 (Biosystems) imaging systems and quantified using Image Studio (v5.2; LI-COR) or ImageJ software (v13.0.6, NIH, USA). Band intensities were normalized to total protein determined by Ponceau staining.

For analysis of TLR9, MyD88, and NF-kB subcellular localization, cytosolic and nuclear fractions were isolated from placental homogenates using a Nuclear Extraction Kit (Abcam) following the manufacturer’s protocol. Protein abundance was quantified as described above and normalized to total protein using Ponceau S staining. Immunoblots and full antibody details are provided in **Supplementary File 1 (Table S4)** and **Supplementary File 2**.

### Statistical analysis

Data distributions were assessed using the Shapiro-Wilk test. Outliers identified by ROUT (Robust regression and Outlier, Q=1%) were excluded prior to hypothesis testing. Data from placentas from female and male fetuses were analyzed using two-way ANOVA followed by Sidak’s multiple comparisons test. For other comparisons, one-way ANOVA was used for normally distributed data with equal variances, followed by Tukey’s multiple comparisons test. When assumptions of equal variance were violated, Welch/Brown-Forsythe ANOVA was used with appropriate post hoc tests (e.g., Dunnett’s T3). For data that were not normally distributed, Kruskal-Wallis tests were used, followed by Dunn’s multiple comparisons. Data are presented as mean ± standard deviation (SD), unless otherwise indicated. For RT-qPCR, statistical analyses were performed on ΔCt values. The significance level was set to α = 0.05. Statistical analyses for placenta and live neonate investigations were conducted in GraphPad Prism (v11;GraphPad Software, San Diego, CA, USA). Exact p-values are reported for all analyses. For multi-endpoint datasets, full ANOVA output (F statistics, degrees of freedom, and exact p-values for main effect and interactions) is provided in supplementary materials.

Stillbirth outcomes were analyzed at the dam (litter) level using Poisson regression, with the number of stillborn pups per litter as the dependent variable. Treatment group was the main independent variable, and total litter size was included as a covariate to account for the increased likelihood of stillbirth events in larger litters. Poisson regression was selected given the count nature of the outcome. The small sample size precluded models better suited for high rates of zeros (e.g., negative binomial regression, zero-inflated regression) (35). The constant variance assumption was evaluated by comparing mean and variance within each treatment group. Model fit was assessed using Pearson goodness-of-fit to evaluate residuals for overdispersion. A χ^2^ likelihood ratio test was used to determine whether inclusion of treatment group significantly improved model fit relative to a null model containing litter size only. Treatment effects are expressed as incidence rate ratios (IRR) with 95% confidence intervals. Stillborn analysis was conducted in R (version 4.4.1) (36).

## Results

### Study 1: Specificity of mtDNA-induced inflammation

A 4-hr exposure to mtDNA, but not nDNA, increased expression of *il1b*, *tnfa*, *il10* mRNA (all Welch’s ANOVA: *il1b*, F(2, 22) = 92.5, p < 0.0001; *tnfa,* F(2, 9.8) = 18.8, p = 0.0004; *il10,* F(2, 15.1) = 5.3, p = 0.02; **Figure 2A-C**). Dunnett’s T3 post hoc test showed that placentas from mtDNA-treated rats had greater expression of *il1b*, *tnfa*, *and il10* than saline controls (p ≤ 0.027), whereas placentas from nDNA-treated rats did not differ from saline-treated controls (p ≥0.154). These data demonstrate that mtDNA uniquely induces an inflammatory transcriptional response in the placenta 4 h after exposure.

**Figure 2.**
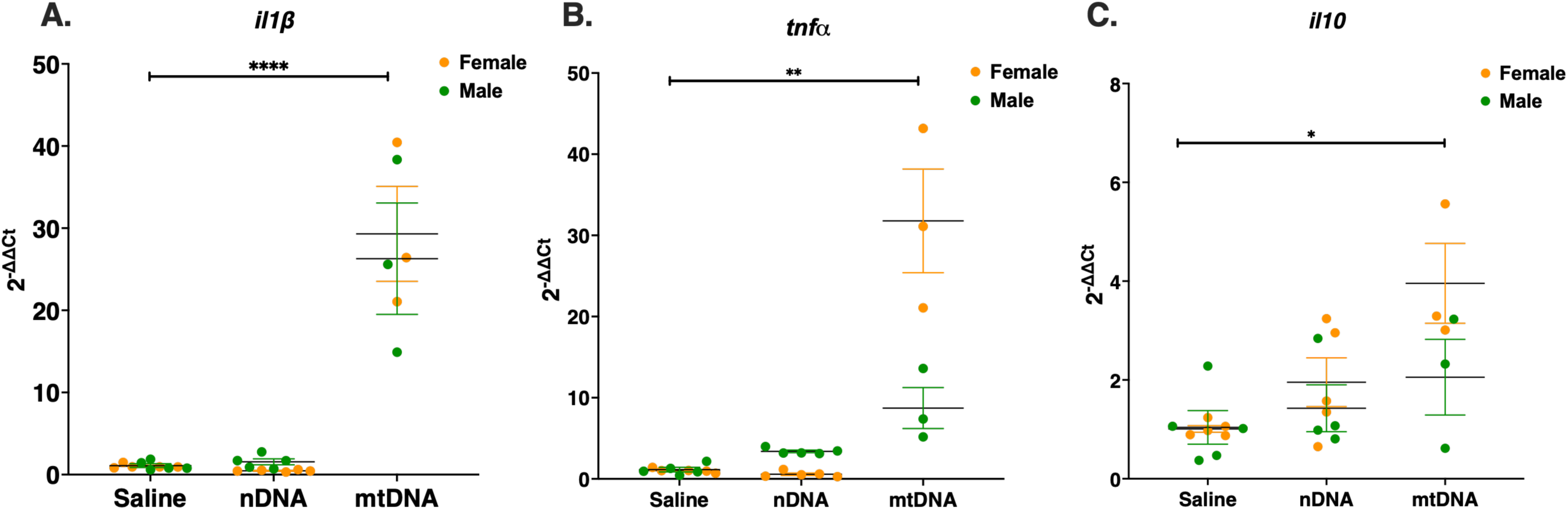
Cytokine mRNA expression in rat placentas four hours after exposure to purified mtDNA. Relative mRNA expression (2^-ΔΔCt^) of (A) interleukin-1β (*il1β*), (B) tumor necrosis factor alpha (*tnfα*), and (C) interleukin-10 (*il10*) in placental tissues from pregnant rats treated with saline, nuclear DNA (nDNA), or mitochondrial DNA (mtDNA) extracted from rat liver. Statistics were performed on ΔCt values using Welch ANOVA followed by Dunnett’s T3 multiple comparisons test. Data are presented as means ± SD. Symbols indicate sex of associated fetus (female, orange; male, green). Saline, n = 10 placentas (5 females, 5 males); nDNA, n = 10 placentas (5 females, 5 males); mtDNA, n = 6 placentas (3 females, 3 males), *p<0.05, **p<0.01, ***p<0.001, ****p<0.0001.

To determine whether these placental responses were accompanied by systemic changes, we assessed ccf-mtDNA and plasma cytokines 4 h after exposure. At this time point, there were no differences in ccf-mtDNA copy number between saline-treated and mtDNA-treated rats, consistent with rapid clearance of circulating DNA (**Supplementary File 1: Figure S1**). Plasma cytokine concentrations at 4 h were largely unchanged by mtDNA exposure, with no group differences in TNF-α and IL-6. Interestingly, MCP-1 concentrations were lower in the mtDNA-treated group compared to the saline-treated controls (**Supplementary File 1: Figure S1**).

We observed a visual divergence in **Figures 2B-C** between placentas from male and female fetuses. Because Study 1 was designed to determine whether mtDNA induces placental inflammation, rather than to evaluate the influence of fetal sex, the study was not powered for sex-specific analyses. Nevertheless, this observation generated the hypothesis that fetal sex may modify placental responses to mtDNA exposure. Subsequent experiments were therefore specifically designed to examine the influence of fetal sex on placental inflammatory signaling.

### Study 2: Sex-dependent placental inflammatory responses to mtDNA

To provide a broad assessment of the transcriptional inflammatory response to purified mtDNA 4-hr after exposure, we measured a panel of pro-inflammatory *(il1β, il6, tnfα)*, anti-inflammatory *(il4, il10),* T cell-associated *(ifnγ)* cytokine genes, as well as mRNA of chemokine/immune cell markers *(mcp1, f4/80, mmp8)*.

There was a significant treatment-by-sex interaction for *il6*, *il1b*, *ifnγ*, *il4*, and *mcp1* (two-way ANOVA; **Figure 3A-E, Supplementary File 1: Table S5**). Post hoc testing showed that mtDNA increased *il6* and *il1β* mRNA in male placentas (p ≤ 0.0004), but not in female placentas (p ≥ 0.075). Conversely, mtDNA increased *ifnγ* in female placentas (p = 0.0004) but not in male placentas (p = 0.997). *il4* and *mcp1* increased in both sexes (p ≤ 0.0015), with a greater response in females **(Figure 3D-E).** For *tnf*α, *il10*, and *f4/80,* we observed main effects of sex and treatment, without significant interactions (**Figure 3F-H, Supplementary File 1: Table S5**). There were no main effects of sex or treatment and no interaction for *mmp8* **(Supplementary File 1: Table S5)**. These data demonstrate that mtDNA elicited a distinct inflammatory response between placentas from male and female fetuses.

**Figure 3.**
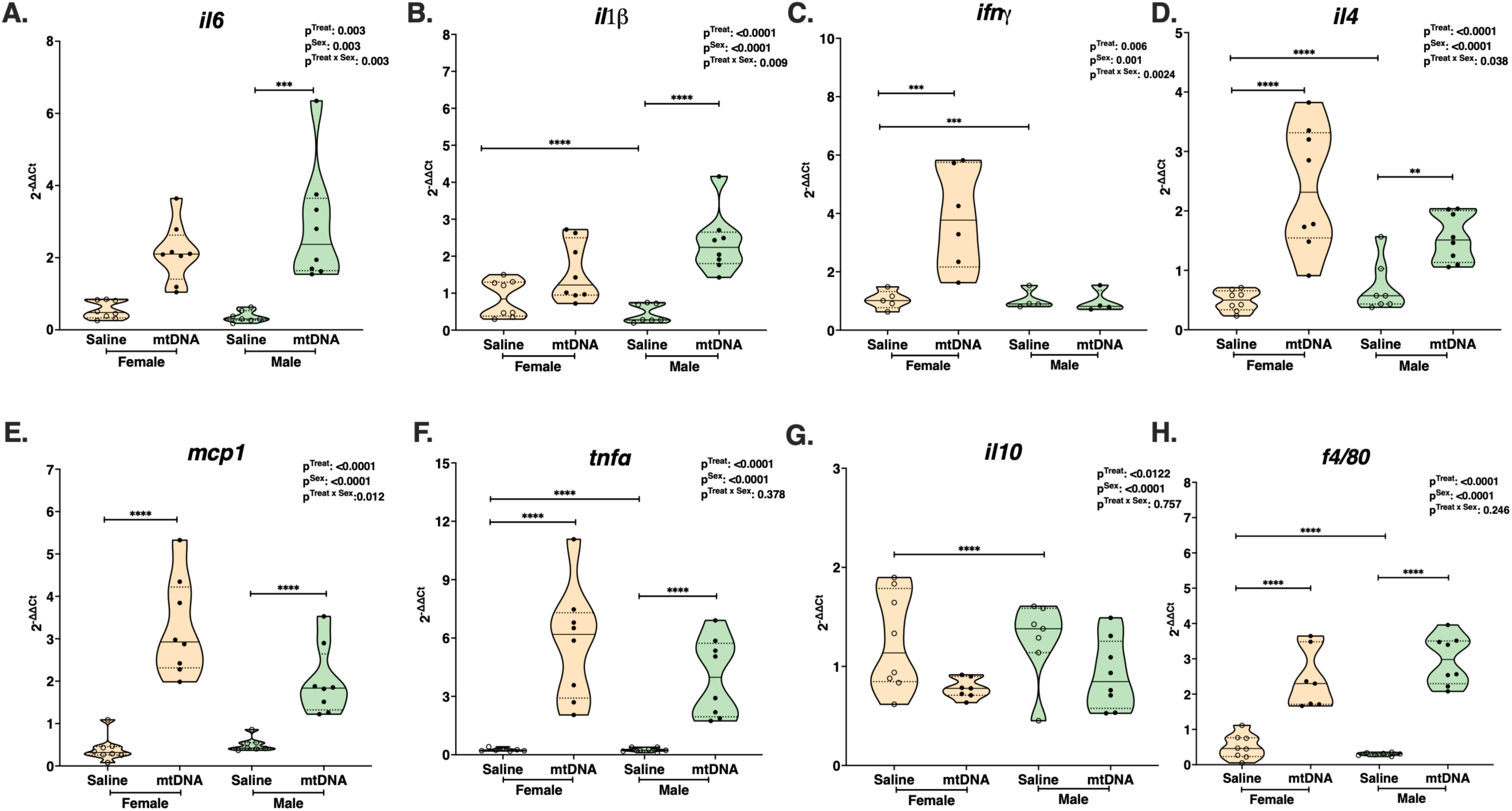
Cytokine mRNA expression in rat placentas from male and female fetuses four hours after exposure to purified mtDNA. Relative mRNA expression (2^-ΔΔCt^) of (A) interleukin-6 (*il6*), (B) interleukin-1β (*il1β*), (C) interferon γ *(ifnγ*), (D) interleukin-4 (*il4*), (E) monocyte chemoattractant protein-1 *(mcp1)*, (F) tumor necrosis factor alpha (*tnfα*), (G) *il10,* and (H) *f4/80* in placentas from female (orange) and male (green) fetuses from pregnant rats treated with saline (open circles) or mitochondrial DNA (mtDNA, closed circles). Statistics were performed on ΔCt values using two-way ANOVA followed by Šidák’s multiple comparisons test. Truncated violin plots show individual placentas. One male and one female placenta were analyzed from each litter. Sample sizes (placentas) were: *il6*, *il1β, il4, mcp1:* female saline, n = 8; female mtDNA, n = 8; male saline, n = 7; male mtDNA, n = 8. For *tnfα, il10, and f4/80,* n = 7-8 placentas per sex-treatment group (one outlier removed). For *ifnγ,* n = 4-5 placentas per sex-treatment group. Solid lines indicate medians and dotted lines indicate quartiles. *p<0.05, **p<0.01, ***p<0.001, ****p<0.0001.

At baseline (saline-treated groups), female placentas had higher expression of *il1β*, *tnfα*, *ifnγ, f4/80*, and *il10* compared with male placentas (p ≤ 0.0003), whereas male placentas showed greater *il4* expression (p < 0.0001). These findings indicate sexual dimorphism in the baseline transcriptional profiles of pro– and anti-inflammatory cytokines.

Placental protein abundance of IL-1β, TNF-α, IL-4, and IL-10 was not altered 4 h after exposure to mtDNA **(Supplementary File 1: Table S6)**. There was a treatment effect for macrophage colony-stimulating factor (M-CSF) and regulated upon activation, normal T cell expressed and secreted (RANTES) protein content (Two-way ANOVA, treatment effect: p=0.0014 and p=0.0396, respectively). These effects were driven primarily by a reduction of M-CSF and RANTES expression in female placentas from baseline levels **(Supplementary File 1: Table S6)**. These findings indicate that 4 h of exposure to mtDNA were associated with a sex-dependent reduction in signals associated with monocyte/macrophage recruitment, but no effects on cytokine protein content.

### Study 3: TLR9 involvement in inflammatory responses to mtDNA

A 4-h exposure to extracellular mtDNA increased *tlr9* mRNA in both female and male placentas without a treatment-by-sex interaction (Treatment, F(1, 27) = 31.34, p < 0.0001, **Figure 4A**). There was also a sex effect, with female placentas having higher baseline expression (saline-treated groups) than male placentas (F(1, 27) = 218.8, p < 0.0001, **Figure 4A**). Cytosolic TLR9 protein abundance was increased in response to mtDNA in female but not in male placentas **(Figure 4B)**. Specifically, we observed a significant treatment-by-sex interaction (F(1, 23) = 5.661, p = 0.026) and main effects for sex and treatment (Sex, F(1, 23) = 5.661, p = 0.026; Treatment, F(1, 23) = 5.338, p = 0.03). mtDNA increased TLR9 protein abundance in female placentas (Sidak’s post hoc test, p = 0.007) but not in male placentas (Sidak’s post hoc test, p > 0.999; **Figure 4B**). We noted a similar pattern for placental abundance of MyD88, which is downstream of TLR9 activation (Treatment-by-sex interaction: F(1, 19) = 14.33, p = 0.001; Sex: F(1, 19) = 14.33, p = 0.001; Treatment, F(1, 19) = 4.838, p = 0.04, **Figure 4C**). Exposure to mtDNA increased MyD88 protein abundance in female placentas (Sidak’s post hoc test, p = 0.001) but not in male placentas (Sidak’s post hoc test, p = 0.637). Together, these data suggest that exposure to mtDNA activates components of TLR9 signaling in female placentas, while this activation is less robust in male placentas at 4 h after exposure to purified mtDNA.

**Figure 4.**
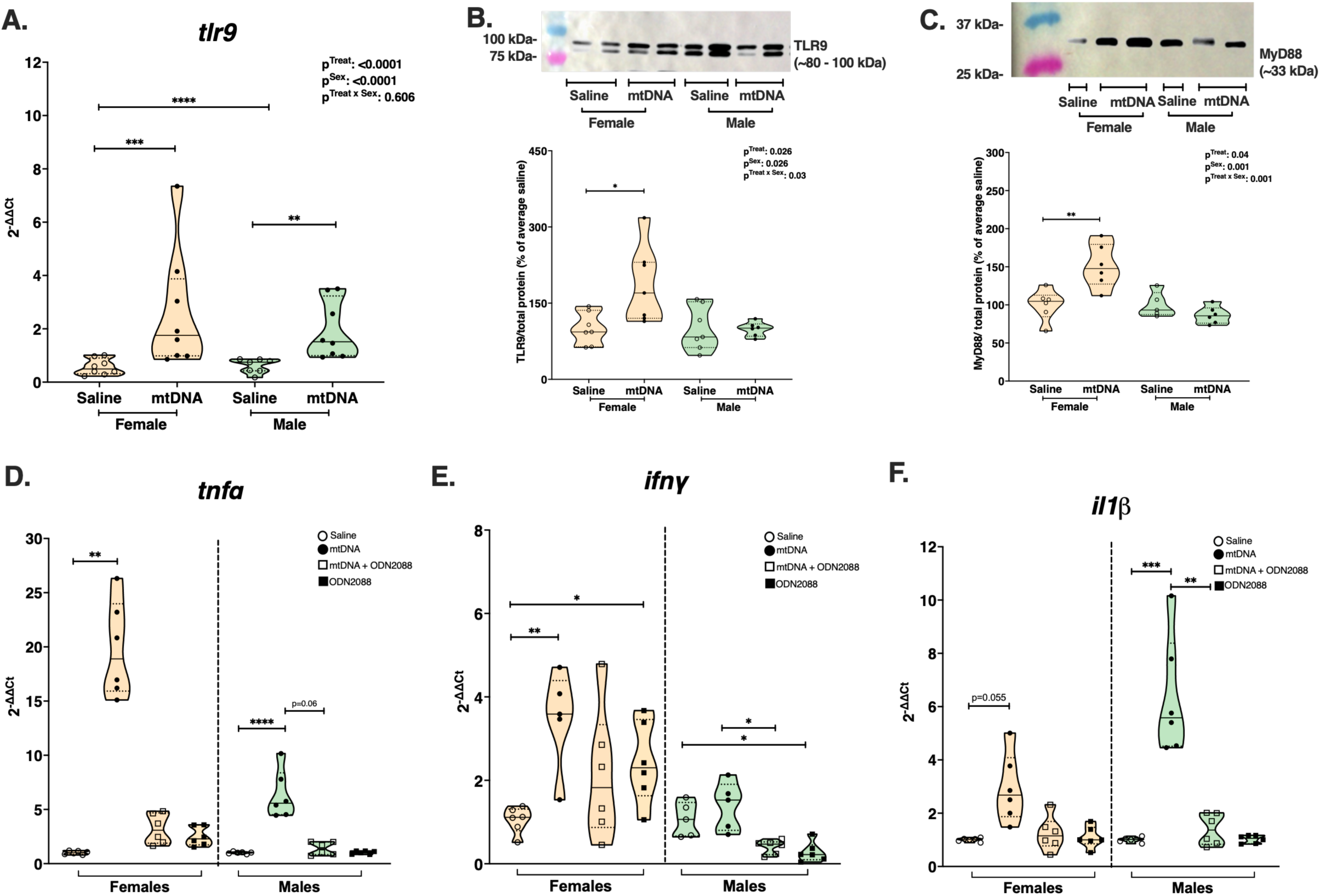
Toll-like receptor 9 (TLR9) involvement in inflammatory responses to mtDNA four hours after exposure. (A) Relative *tlr9* mRNA expression, (B) TLR9 protein expression, (C) MyD88 protein expression, and relative mRNA expression (2^-ΔΔCt^) of (D) tumor necrosis factor alpha *(tnfα)*, (E) interferon γ *(ifnγ)*, (F) interleukin-1β *(il1β)* in placentas from female (orange) and male (green) fetuses from pregnant rats treated with saline (open circles), mitochondrial DNA (mtDNA, closed circles), mtDNA and toll-like receptor 9 (TLR9) antagonist ODN2088 (open squares), or ODN2088 alone (closed squares). Protein levels were normalized to total protein and expressed relative to saline-treated controls (set to 100%). For all analyses, truncated violin plots show individual placentas. One male and one female placenta were analyzed from each litter. Solid lines indicate medians and dotted lines indicate quartiles. (A-C) Statistics were performed using two-way ANOVA followed by Šidák’s multiple comparisons test. Sample sizes were *tlr9,* n = 7-8 placentas per sex-treatment group; TLR9, n = 6-7 placentas per sex-treatment group; MyD88, n = 5-6 placentas per sex-treatment group. (D-F) Statistics were performed separately within each sex group. One-way ANOVA followed by Dunnett’s multiple comparisons was used for *ifnγ* (n = 5-6 placentas per treatment group within each sex). Kruskal-Wallis test followed by Dunn’s multiple comparisons was used for *tnfα* in female placentas (n = 6 placentas per treatment group). Welch ANOVA followed by Dunnett’s T3 multiple comparisons was used for *il1β* in male and female placentas (n = 6 placentas per treatment group) and *tnfα* in male placentas (n = 6 placentas per treatment group). *p<0.05, **p<0.01, ***p<0.001, ****p<0.0001.

We then pharmacologically tested TLR9 requirement in placental inflammatory responses to mtDNA. We treated dams with mtDNA (or saline) with and without a TLR9 antagonist (ODN2088) and evaluated whether ODN2088 modified the mtDNA response by comparing mtDNA vs. mtDNA + ODN2088 within each sex, and we also assessed the effect of ODN2088 alone relative to saline. In female placentas, mtDNA increased mRNA expression of *tnfα* and *ifnγ* (p = 0.0021 and p = 0.0037, respectively) and ODN2088 did not reduce mtDNA-associated increases for the cytokines examined (all p ≥ 0.2375; **Figure 4D-F**). In male placentas, mtDNA increased *tnfα* and *il1β* mRNA (p < 0.0001 and p = 0.0006, respectively). ODN2088 prevented the mtDNA-associated increase in *il1β* (p = 0.0048, **Figure 4F**), with a trend toward reduction for *tnfα* (p =0.061). Interestingly, ODN2088 altered basal *ifnγ* expression in both female and male placentas (p = 0.019 and p = 0.015, respectively). Together, these findings provide evidence that ODN2088 prevented the mtDNA-associated *il1β* increase and tended to reduce *tnfα* in male placentas but had no effect in the cytokine examined in female placentas at 4 h after exposure to mtDNA.

To assess NF-κB activation, we examined the nuclear:cytosolic ratio of the p65 subunit, which reflects net translocation from cytosol to nucleus. In female placentas, neither mtDNA nor ODN2088 affected the nuclear:cytosolic p65 ratio (p ≥ 0.21, **Figure 5A-B**), despite mtDNA increasing the abundance of both cytosolic and nuclear p65 individually (p = 0.026 and p = 0.0058, respectively; **Supplementary File 1: Figure S2**). ODN2088 reduced these individual fraction increases (mtDNA vs. mtDNA+ODN2088: cytosolic p65, p = 0.049; nuclear p65, p = 0.039; **Supplementary File 1: Figure S2**), yet the ratio remained unchanged. In male placentas, neither mtDNA nor ODN2088 affected cytosolic p65 (p ≥ 0.09, **Supplementary File 1: Figure S2**), nuclear p65 (p ≥ 0.7, **Supplementary File 1: Figure S2**) or their ratio (p ≥ 0.64; **Figure 5A-B**).

**Figure 5.**
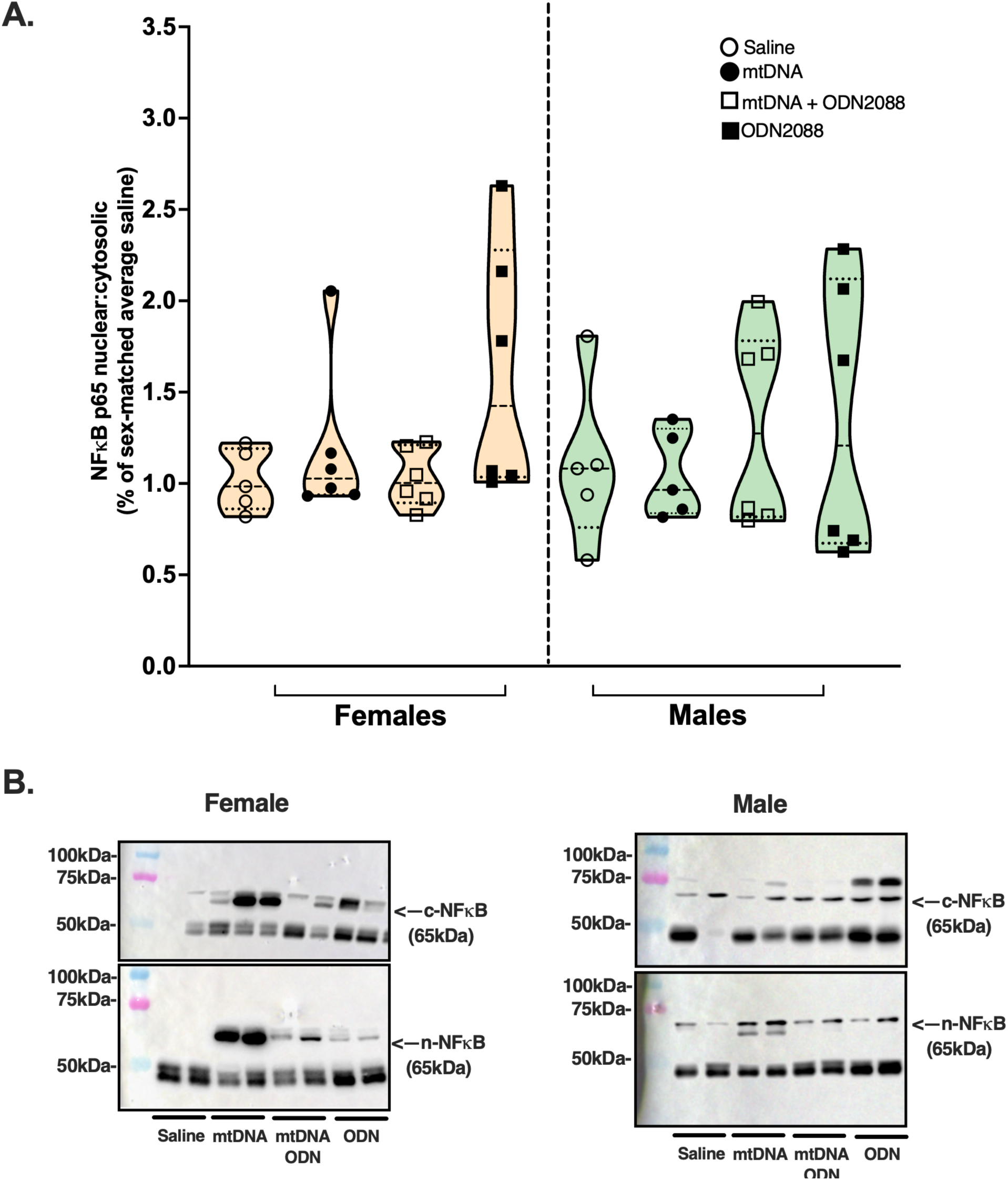
Nuclear factor kappa-light-chain-enhancer of activated B cells (NF-κB) p65 nuclear translocation in placentas from male and female fetuses four hours after in vivo pharmacological antagonism of Toll-like receptor 9 (TLR9). (A) Nuclear:cytosolic ratio of NF-κB p65 subunit protein abundance, reflecting net translocation from cytosol to nucleus, in placentas from female (orange) and male (green) fetuses of pregnant rats treated with saline (open circles), mitochondrial DNA (mtDNA; closed circles), mtDNA + TLR9 antagonist ODN2088 (open squares), or ODN2088 alone (closed squares). Protein levels in cytosolic and nuclear fractions were independently normalized to total protein before ratio calculation. (B) Representative western blot images of NF-κB p65 in cytosolic and nuclear fractions from placentas of female (top) and male (bottom) fetuses. Cytosolic and nuclear p65 protein abundances are shown separately in Supplementary Figure S2. Statistics were performed using Welch ANOVA followed by Dunnett’s T3 multiple comparisons test. Truncated violin plots show individual placentas. One male and one female placenta were analyzed from each litter. Sample sizes were n = 5-6 placentas per sex-treatment group. Solid lines indicate medians and dotted lines indicate quartiles. *p<0.05, **p<0.01, ***p<0.001, ****p<0.0001.

Taken together, these findings indicate that TLR9 signaling contributes to mtDNA-induced placental inflammation in a sex-dependent manner. In male placentas, ODN2088 prevented the mtDNA-associated *il1β* increase and tended to reduce *tnfα* providing direct pharmacological evidence for TLR9 involvement. In female placentas, ODN2088 reduced mtDNA-associated increases in cytosolic and nuclear p65, yet cytokine mRNA responses were unaffected and the nuclear:cytosolic ratio remained unchanged in both sexes.

### Study 4: Inflammation resolution and antioxidant defense mechanisms

To determine whether the placental inflammatory responses to extracellular mtDNA persist, we measured expression of selected cytokines in a separate cohort 24 h post-treatment, focusing on cytokines that showed sex-specific patterns in earlier time points (i.e., 4 h post-treatment). We found no treatment effect (p ≥ 0.105) and no treatment-by-sex interaction (p ≥ 0.085) in expression of *il6, il1β,* and *il4* **(Supplementary File 1: Figure S3).** There was a main effect of sex for *il6* and *il1β* (p ≤ 0.024).

Given the absence of a treatment effect on these cytokines at 24 h, we next evaluated whether downstream oxidative stress-related markers were altered. At 24 h post-treatment, *sod2 and catalase* expression showed sex-specific patterns that differed from the cytokine resolution observed at this timepoint (Treatment-by-sex interaction, p ≤ 0.0043; **Figure 6E-F, Supplementary File 1: Table S7**). Specifically, *sod2* and *catalase* were lower in male placentas from saline-treated dams compared with mtDNA-treated dams (p ≤ 0.0184), whereas these enzymes were unchanged in female placentas (p ≥ 0.073).

**Figure 6.**
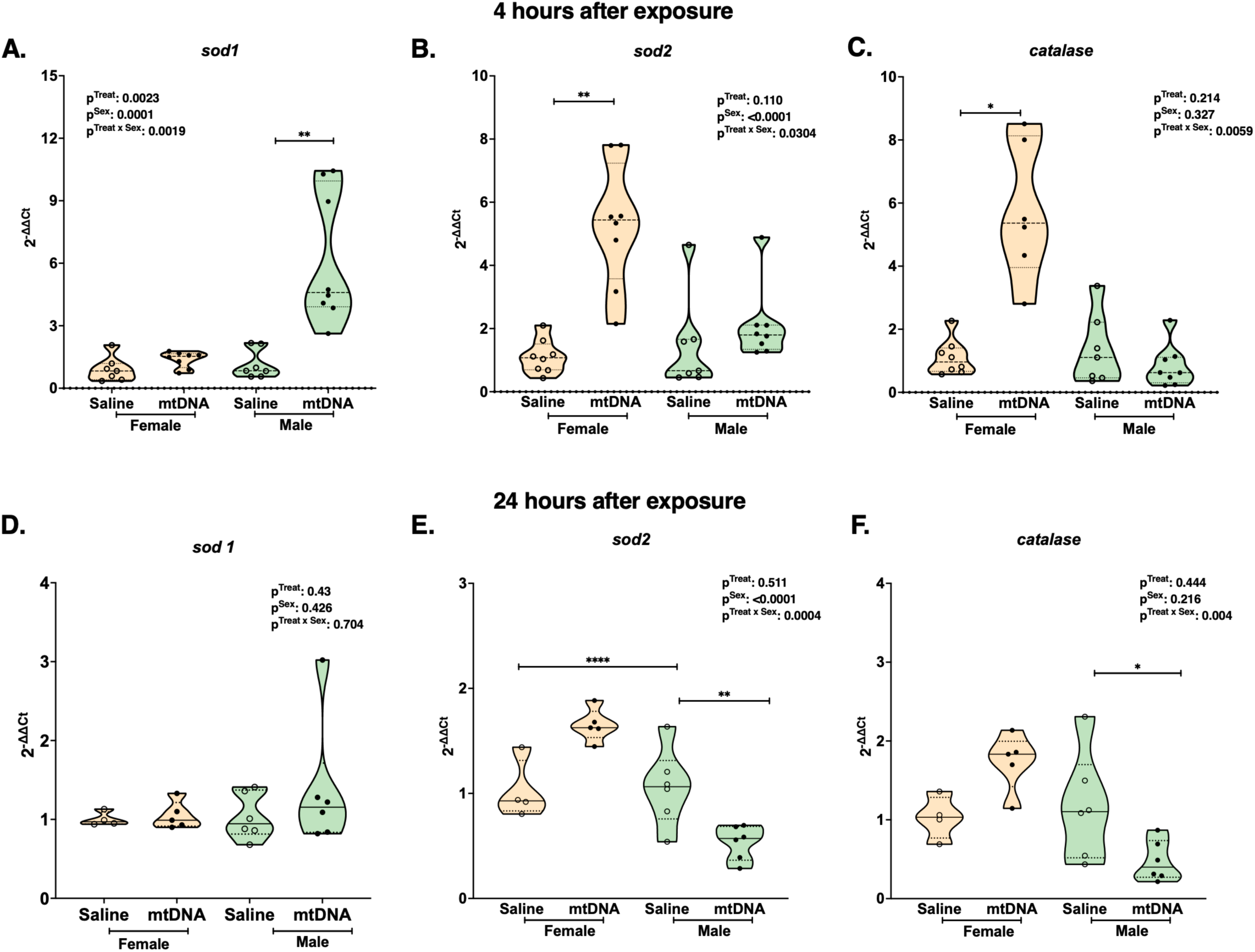
Placental antioxidant enzyme mRNA expression 4 h and 24 h after in vivo exposure to purified mtDNA. Relative mRNA expression of antioxidant enzymes was quantified by qRT-PCR in placentas from female (orange) and male (green) fetuses of pregnant rats treated with saline (open circles) or mitochondrial DNA (mtDNA; closed circles). Top panels show responses 4 h post-treatment: (A) *sod1*, (B) *sod2*, and (C) *catalase*. Bottom panels show responses 24 h post-treatment: (D) *sod1*, (E) *sod2*, and (F) *catalase*. Statistics were performed on ΔCt values using two-way ANOVA followed by Šidák’s multiple comparisons test. Truncated violin plots show individual placentas. One male and one female placenta were analyzed from each litter when available. Sample sizes were n = 7-8 placentas per sex-treatment group for the 4 h time point and n = 4-6 placentas per sex-treatment group for the 24 h time point. Solid lines indicate medians and dotted lines indicate quartiles. p<0.05, **p<0.01, ***p<0.001, ****p<0.0001.

These findings prompted us to examine whether sex-specific differences in antioxidant enzyme expression were already present at 4 h post-treatment. Analysis of placental tissues collected in Study 2 (described above) revealed that at 4 h post-treatment, mtDNA altered placental antioxidant enzyme expression in a fetal sex-dependent manner **(Figure 6A-C, Supplementary File 1: Table S7)**. Specifically, *sod1* increased in male (p < 0.0001) but not in female placentas (p = 0.87), whereas *sod2* and *catalase* increased in female (p ≤ 0.03) but not in male placentas (p ≥ 0.115).

Together, these data suggest that mtDNA exposure is associated with a sex-specific shift in antioxidant enzyme expression that is detectable as early as 4 h and persists through 24 h post-treatment **(Figure 6A-F)**, whereas the mtDNA-associated inflammatory transcriptional responses observed at 4 h are transient and no longer detectable at 24 h.

### Study 5: Pregnancy and neonatal outcomes after exposure to mtDNA during pregnancy

Exposure to extracellular mtDNA during pregnancy did not affect gestational length (median (IQR), saline: 22 (0), mtDNA: 22 (1), Mann-Whitney *U* test, p = 0.2), or neonatal body weight, crown-rump length, and abdominal girth measured within 12 h after delivery **(Supplementary File 1: Figure S4)**. Treatment effects on stillbirth counts per litter were analyzed using Poisson regression, adjusting for total pups per litter **(Table 1).** Treatment was nearly significantly associated with stillbirth count (p = 0.069), with mtDNA-treated dams exhibiting an approximately 75% higher expected stillbirth count than saline-treated dams (estimated marginal means: mtDNA, 1.03 ± 0.40 vs. saline, 0.24 ± 0.17; IRR = 4.23, 95% CI [0.89, 20.1], **Figure 7**). Litter size was not significantly associated with stillbirth count in the full model (p = 0.34). Inclusion of treatment significantly improved model fit over a litter-size-only null model (likelihood ratio test, χ^2^ = 4.13, p = 0.042; **Supplementary File 1: Table S8**). Collectively, our data show that extracellular mtDNA exposure during pregnancy did not alter gestational timing or neonatal biometrics, but it was associated with a higher estimated expected stillbirth count per litter.

**Figure 7.**
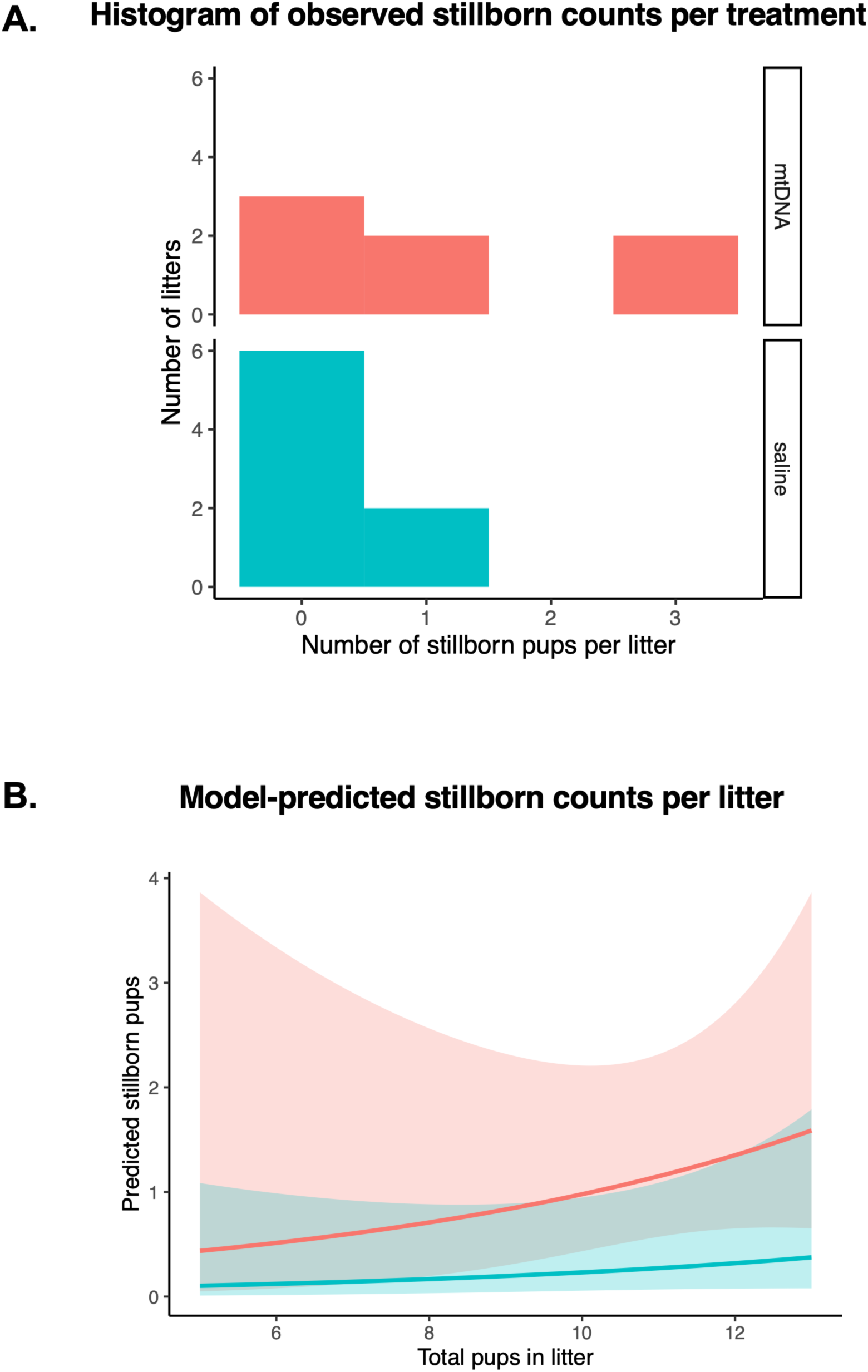
Stillbirth outcomes after gestational exposure to purified mtDNA: observed distribution and Poisson regression estimates. (A) Histogram of the number of stillborn pups per litter in saline– and mtDNA-treated dams (bin size = 1). Each litter represents one observation (saline n = 8; mtDNA n = 7). (B) Predicted number of stillborn pups as a function of total litter size from a Poisson regression model. Curves represent model-predicted stillbirth counts across a range of litter sizes, and shaded ribbons indicate 95% confidence intervals. Predictions are shown for saline-treated (teal) and mtDNA-treated (coral) dams. Model predictions were generated from 100 simulated litter sizes per treatment group (N = 200).

**Table 1.**
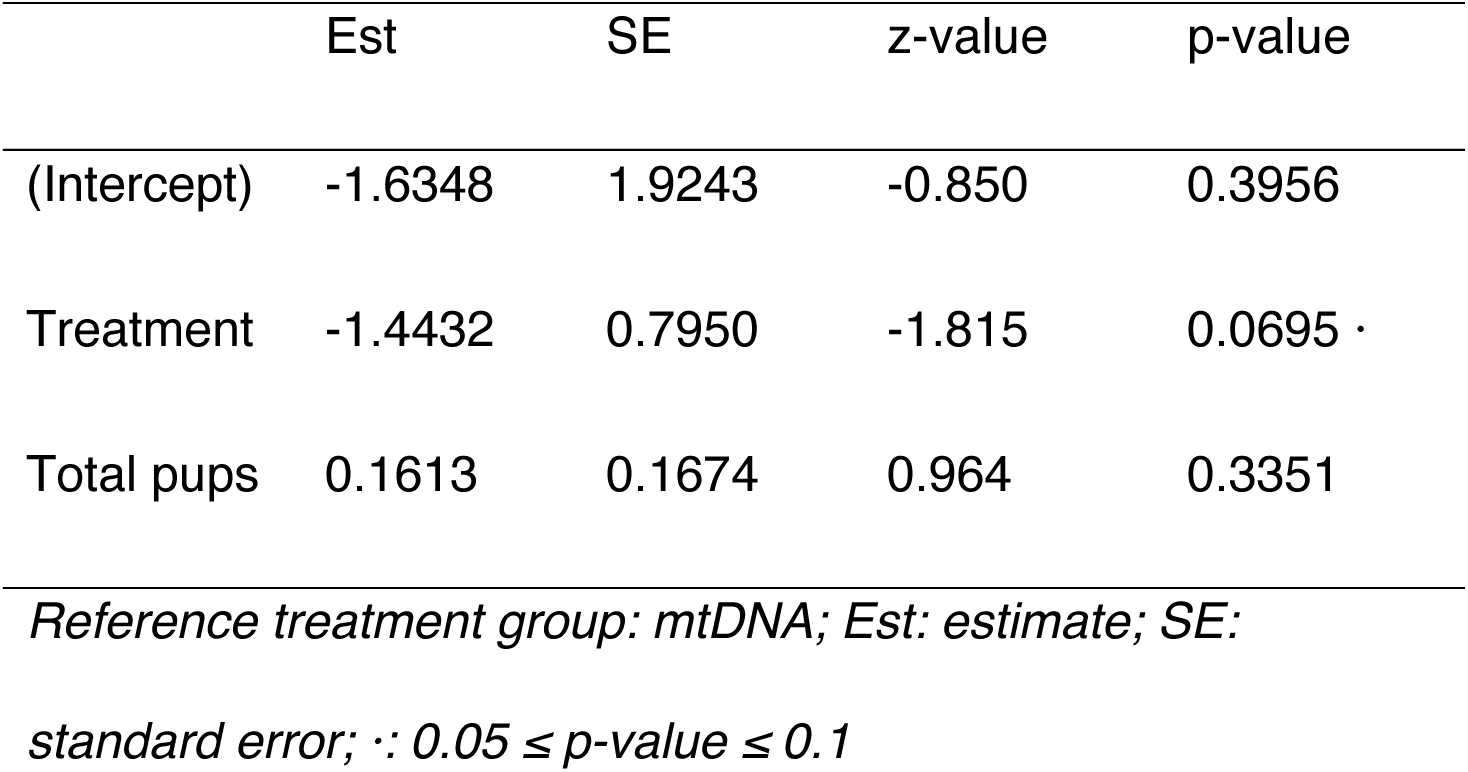
Simple Poisson regression results. Comparison of stillborn pups as a function of total litter size by treatment.

|  | Est | SE | z-value | p-value |
| --- | --- | --- | --- | --- |
| (Intercept) | -1.6348 | 1.9243 | -0.850 | 0.3956 |
| Treatment | -1.4432 | 0.7950 | -1.815 | 0.0695 |
| Total pups | 0.1613 | 0.1674 | 0.964 | 0.3351 |

*Reference treatment group: mtDNA; Est: estimate; SE:*
*standard error; $\therefore 0.05 \leq p\text{-value} \leq 0.1$*

## Discussion

The placenta is an immunologically active organ whose inflammatory responses to circulating danger signals may shape both fetal development and maternal health. Among these signals, extracellular mtDNA has emerged as a candidate DAMP capable of activating innate immune receptors (5, 37). Whether and how mtDNA drives placental inflammation and whether fetal sex modifies this response was poorly understood. The present study identifies extracellular mtDNA as a sex-differentiated placental inflammatory stimulus with implications for understanding how circulating maternal danger signals may contribute to placental and fetal vulnerability in complicated pregnancies.

Pregnancies complicated by inflammation and adverse outcomes, including preeclampsia, are associated with distinct ccf-mtDNA profiles compared to uncomplicated pregnancies at various gestational stages (7–10). Deviations from its normal gestational profile likely reflect dysregulation of one or more mechanisms governing mtDNA release, clearance, or tissue uptake (38–40). The present study did not aim to resolve these mechanisms but instead addressed a more focused question: whether extracellular mtDNA is sufficient to elicit a placental inflammatory response.

Consistent with previously reported rapid clearance of extracellular DNA from the circulation (38, 41), ccf-mtDNA concentrations did not differ between mtDNA-treated and saline-treated dams at 4 h post-exposure, indicating that the administered mtDNA was no longer detectable in maternal plasma at this timepoint. These findings indicate that the placental inflammatory responses observed in mtDNA-treated animals reflect the biological consequences of transient innate immune activation rather than persistent supraphysiological circulating mtDNA levels and support the physiological relevance of this experimental model. Despite absence of increased circulating mtDNA at 4 h, placental inflammatory transcription was robustly increased. Importantly, these responses were specific to mtDNA, as intravenous administration of nDNA at an equivalent dose did not elicit detectable placental inflammatory responses, establishing the molecular specificity of the effect.

The pattern of placental cytokine induction suggests that mtDNA engages shared inflammatory pathways in both sexes, while preferentially engaging *il1β/IL6-associated* pathways in male placentas and *ifnγ*-associated pathways in female placentas. Male-biased *il1β/il6* responses may reflect preferential activation of inflammasome-associated (42) and monocyte/macrophage-linked pathways involved in inflammatory tissue remodeling (43). In contrast, the selective induction of *ifnγ* in female placentas suggests an interferon-associated immune phenotype, potentially involving natural killer cell– or T cell-associated immune programs that are well characterized at the maternal-fetal interface (43–45). These transcriptional sex differences appeared within 4 h of mtDNA exposure before corresponding changes in cytokine protein abundance were detected, suggesting that early sex-specific inflammatory programming precedes measurable cytokine accumulation.

In contrast, mtDNA altered placental M-CSF and RANTES protein abundance, particularly in female placentas, suggesting that chemokine-mediated immune recruitment pathways may respond earlier than cytokine protein production. Although not directly examined in the present study, the sex-specific cytokine pattern in response to mtDNA may reflect differences in baseline immune setpoints (22), as female placentas exhibited a more robust basal cytokine profile compared with male placentas. This sexual dimorphism in baseline transcriptional profiles may reflect a faster trajectory of immune system development in females compared to males (22), which would influence responsiveness to immune triggers, including extracellular mtDNA.

Differences in placental immune cell composition and sex chromosome-dependent regulation of inflammatory signaling are potential drivers of sex differences in placental immune responses. A recent study using the four-core genotypes mouse model demonstrated that XX gonadal females exhibit a distinct placental proinflammatory signature largely independent of maternal immune activation, alongside greater placental buffering capacity, whereas gonadal males showed greater susceptibility to maternal immune activation-related impairments in umbilical blood flow (46). Our findings support the concept that both sex chromosome complement and gonadal hormones independently shape placental inflammatory tone and responsiveness, providing a broader experimental framework for the sex-specific placental responses to extracellular mtDNA observed here. IRAK1, a key mediator of MyD88-dependent TLR signaling, is X-linked and escapes X-inactivation, resulting in higher expression in XX compared to XY cells (47), raising the possibility that higher IRAK1 expression in XX cells could amplify innate immune signal transduction downstream of TLR9 in female placentas and contribute to the stronger TLR9/MyD88 activation we observed.

TLR9 overexpression and increased signaling activity have been implicated in adverse pregnancy outcomes in humans and experimental models (12–14). Whether fetal sex modulates TLR9-mediated innate immune activation in the placenta, however, had not been previously examined. Our data indicate that mtDNA activated the TLR9 pathway in both sexes, though this response was more pronounced in female placentas. In addition, pharmacological antagonism of TLR9 modified selected mtDNA-associated inflammatory responses in a sex-specific manner. Nevertheless, the partial dependence of placental inflammatory responses on TLR9 signaling, particularly the persistence of cytokine transcriptional responses in female placenta despite antagonism, suggest that additional innate immune sensing pathways contribute to mtDNA-induced inflammation. Extracellular mtDNA can also activate the NLRP3 inflammasome and cytosolic DNA sensors including cGAS-STING (37), neither of which were examined here. Inflammasome activation is particularly consistent with the male-biased *il1β* response we observed, as IL-1β is a canonical inflammasome substrate processed by caspase-1 (48).

Upon CpG recognition, TLR9 traffics from the endoplasmic reticulum to endolysosomal compartments where it recruits MyD88, activating downstream NF-κB signaling and proinflammatory gene transcription (49). To assess NF-κB activation, we examined the nuclear:cytosolic p65 ratio as the primary index of translocation. In female placentas, mtDNA increased both cytosolic and nuclear p65 abundance, and ODN2088 reduced these responses; yet the nuclear:cytosolic ratio was not significantly altered. In male placentas, neither mtDNA nor ODN2088 affected p65 abundance or its distribution. The absence of a significant ratio change despite increases in both p65 fractions in female placentas may reflect an increase in total p65 protein content rather than selective nuclear translocation. Alternatively, it may indicate that translocation occurred and resolved prior to the 4 h collection window, consistent with reports of NF-κB nuclear translocation within 0.5-6 h of innate immune stimulation (50).

The placental transcriptional response observed at 4 h was not reflected in the plasma cytokine profile at this timepoint, as there were no differences in TNF-α or IL-6 between mtDNA-treated and saline-treated dams. Discrepancies between circulating cytokine levels and immune gene expression has been previously reported and attributed to strength of the stimulus or differences in decay rate and transcriptional regulation networks (15, 51, 52). Notably, circulating MCP-1 concentrations were lower in mtDNA-treated dams compared to saline controls, which may reflect regulatory mechanisms during inflammation, potentially involving altered monocyte trafficking, tissue redistribution, or suppression of further MCP-1 production (53).

The mtDNA-induced placental inflammatory response was robust and rapid, with transcriptional normalization by 24 h after exposure. Assessment of cytokine mRNA expression at this later timepoint revealed no significant treatment effects or treatment-by-sex interactions for *il6*, *il1β*, and *il4*, in contrast to the strong transcriptional responses observed at 4 h. Several mechanisms may contribute to this resolution, including rapid negative feedback regulation of NF-κB signaling regulation (54), receptor desensitization following ligand exposure (55), clearance of mtDNA from the maternal circulation, or inhibitory mechanisms to suppress inflammatory responses and prevent adverse pregnancy outcomes (56, 57). We posit that the absence of detectable transcript changes at 24 h does not indicate lack of lasting biological effects; instead, these data suggest that inflammatory responses at 4 h may have initiated downstream cellular responses, such as immune cell recruitment or oxidative stress.

In agreement with this interpretation, mtDNA was associated with sex-specific changes in placental antioxidant enzyme expression that were detectable at both timepoints, suggesting a transition from acute inflammation to altered redox regulation. Previous studies have shown that pro-inflammatory cytokines such as IL-1β, IL-6, IFN-γ can stimulate reactive oxygen species (ROS) production through mitochondrial dysfunction and activation of oxidative pathways (58). In female placentas, *sod2* and *catalase* were induced at 4 h and returned to baseline by 24 h, suggesting a coupled and temporarily coordinated inflammatory and antioxidant response. In male placentas, *sod1* was transiently induced at 4 h only, while *sod2* and *catalase* did not change at 4 h but decreased relative to saline controls at 24 h, indicating a delayed or dysregulated redox compensation. This sex-specific temporal dissociation between inflammatory and antioxidant responses may reflect distinct cytokine pathways engaged by mtDNA in female vs. male placentas: the male-biased *il1β* response is consistent with sustained oxidative signaling through inflammasome-associated pathways, whereas the female-biased *ifnγ* response may promote earlier and more coordinated antioxidant compensation. These mechanisms remain to be confirmed, however, as they were not directly assessed in the present investigation.

Long-term exposure to extracellular mtDNA during pregnancy did not affect gestational length, neonatal body, crown-rump length, or abdominal girth, suggesting that a single mtDNA challenge at this dose and gestational age does not measurably impair overall fetal growth and placental efficiency. Gross growth metrics, however, may be relatively insensitive to acute placental stress of moderate severity and may not reflect fetal vulnerability. Indeed, our data indicate an association between exposure to mtDNA during pregnancy and stillbirth count, though the relatively small numbers of observations require subsequent confirmatory experiments to draw confident conclusions. We conclude that while the mtDNA challenge did not alter gestational duration or neonatal biometrics, the increase in expected stillbirths suggests that extracellular mtDNA may elevate fetal risk without overtly changing gross growth outcomes.

## Limitations

Several limitations should be considered when interpreting these findings, particularly regarding timing, tissue resolution, and the ability to identify cellular mechanisms. The use of a single dose and defined timing of mtDNA exposure may not fully capture the effects of chronic or repeated exposure scenarios that are more physiologically relevant. Additionally, reliance on whole-placenta homogenates may obscure important cell-type-specific or spatial differences, such as distinctions between maternal and fetal compartments or between the labyrinth and junctional zones.

Discrepancies between transcript and protein levels may reflect differences in timing, assay sensitivity, compartmentalization, or post-transcriptional regulation; however, these processes were not addressed in the present study. Immune characterization was also limited, as direct quantification of immune cell populations was not performed. In the stillbirth analysis, a relatively small sample size, low event counts, and trend-level statistical significance indicate that these findings should be interpreted cautiously and require independent replication. In addition, although litter-level effects were considered, other potential confounders, such as variation in fetal sex ratios within litters, remain important considerations. Finally, because multiple placentas per dam were analyzed, samples within a litter share a common maternal environment and are not fully independent observations. Although mixed models incorporating dam as a random effect would provide more rigorous accounting of this nested structure, the sample sizes in the present studies were insufficient to reliably estimate random effects parameters. Future studies should be prospectively powered and designed to accommodate this analytical approach.

## Conclusion

Collectively, these findings identify extracellular mtDNA as a transient, sex-dependent driver of placental inflammation, acting in part through TLR9 and accompanied by downstream redox alterations, offering molecular insight into why male and female fetuses may follow distinct vulnerability trajectories under inflammatory conditions.

Moving forward, studies that resolve cell-type and spatial specificity will be essential to identify the precise cellular sources and targets of these responses. Expanding investigation into complementary pathways, including inflammasome activation and cGAS-STING signaling, will further refine the mechanisms underlying the observed responses. Equally important will be integrating functional and longitudinal approaches, such as assessments of placental perfusion, vascular reactivity, barrier integrity, and trophoblast stress, to link molecular changes to physiological outcomes. Finally, expanding dose and time-course patterns, incorporating chronic exposure models, and replicating the observed stillbirth signal will be essential to extend these findings.

## Supporting information

Blot images

## Acknowledgments

We thank the Lawrence D. Longo MD, Center for Perinatal Biology Core Facility at Loma Linda University for providing access to equipment and instrumentation necessary to complete the Western blot and RT-qPCR analyses. We acknowledge the staff at the Animal Care Facilities at Loma Linda University for support with animal husbandry and veterinary care.

The authors used Claude (Claude Sonnet 4.6, Anthropic) to assist with grammar, syntax, and clarity editing in portions of this manuscript. All AI-assisted content was reviewed, verified, and approved by the authors, who take full responsibility for the accuracy, integrity, and originality of the published work. Use of this tool was consistent with the ethical policies of the American Physiological Society.

## Data Availability Statement

Raw data used for analyses in this study are available from the authors upon reasonable request.

## Disclaimers

The content is solely the responsibility of the authors and does not necessarily represent the official views of the National Institutes of Health.

## Author Contributions

RNOS and SG conceived and designed research; RNOS, NH, and SG analyzed data; RNOS, NH, SG, LL, IG, GK, MR, NRP, DE performed experiments; RNOS, NH, NRP, CG, and SG interpreted experiments; RNOS and SG prepared figures; RNOS and SG drafted manuscript; all authors edited, revised and approved the final version of the manuscript.

## Figure legends

## Supplementary Materials

**SUPPLEMENTARY FILE 1**:

## Supplementary Tables

- **Supplementary Table S1:** Chemicals and reagents
- **Supplementary Table S2:** Primers for Kdm5c (X chromosome) and kdm5d (Y chromosome) genes used in fetal sex determination
- **Supplementary Table S3:** Primer sequences for quantitative real-time PCR analysis of gene expression in placenta
- **Supplementary Table S4:** Primary and secondary antibodies used for Western blotting
- **Supplementary Table S5:** ANOVA and pos-hoc output for Figure 3A
- **Supplementary Table S6:** Multiplex panel of cytokine and chemokine levels in the placentas from male and female fetuses 4 h after exposure to purified mitochondrial DNA (mtDNA)
- **Supplementary Table S7:** ANOVA and pos-hoc output for Figure 6A-F
- **Supplementary Table S8:** Testing significance of Treatment effect

## Supplementary Figures

- **Supplementary Figure S1:** Circulating cell-free mitochondrial DNA (mtDNA) copy number and plasma concentrations of TNF-α, IL-6, and MCP-1
- **Supplementary Figure S2:** Nuclear factor kappa-light-chain-enhancer of activated B cells (NF-κB) expression in placentas from male and female fetuses
- **Supplementary Figure S3:** Cytokine mRNA expression in rat placentas 24 h after exposure to purified mtDNA
- **Supplementary Figure S4:** Biometrics of neonates from pregnancies exposed to an acute mtDNA challenge during pregnancy

**SUPPLEMENTARY FILE 2:** PCR gel images for sex determination and Western blot images

