## Supplementary Tables & Figures for "Fetal sex shapes placental inflammatory responses to extracellular mitochondrial DNA"

**SUPPLEMENTARY FILE 1: SUPPLEMENTARY TABLES & FIGURES**

### Fetal sex shapes placental inflammatory responses to extracellular mitochondrial DNA

Reneé de Nazaré Oliveira da Silva[1], Nataliia Hula[1], Desirae Escalera[1], Leslie Lopez [1], Gabrielle Kelly[1], Isabelle K. Gorham[2], Megan Rowe[2], Contessa A. Ricci[3], Ciprian Gheorghe[1,4], Nicole R. Phillips[2], Styliani Goulopoulou[1, 4]

[1]Lawrence D. Longo, MD Center for Perinatal Biology, Department of Basic Sciences, Loma Linda University, Loma Linda, California, USA; [2]Department of Microbiology, Immunology, and Genetics, University of North Texas Health Science Center, Fort Worth, Texas, USA; [3]Institute for Research and Education to Advance Community Health (IREACH), College of Nursing, Elson S. Floyd College of Medicine, Washington State University, USA; [4]Department of Gynecology and Obstetrics, Loma Linda University, Loma Linda, California, USA

**Table S1. Chemicals and reagents.**

| Chemical / Reagent Name | Catalog No. | Vendor/City, State, Country |
|---|---|---|
| $\beta$-mercapoethanol | M3148-25 ML | Sigma-Aldrich |
| Bovine Serum Albumin | A6003-25G | Sigma Aldrich |
| Chromogenic Endotoxin Quant Kit | A39552 | Thermo Fisher Scientific, Waltham, MA, USA |
| DNeasy Blood & Tissue Kit | 69504 | Qiagen, Redwood City, CA, USA |
| DreamTaq PCR Master Mixes (2X) | K1081 | Thermo Fisher Scientific |
| Extract-N-AMP-Tissue PCR Kit | XNAT2-1KT | Sigma-Aldrich |
| iQ SYBR Green Supermix | 1708882 | Bio-Rad, Hercules, CA, USA |
| Isoflurane | 200-237 V1 502017 | Vet One Fluriso, Union City, CA, USA |
| miRNeasy Mini Kit | 217004 | Qiagen |
| Mitochondria Isolation Kit | 89801 | Thermo Fisher Scientific |
| Neutralization Solution B | N3910-24ML | Sigma Aldrich |
| Nitrocellulose membrane, 0.45um 30 cm x 3.5 m | 162-0115 | Bio-Rad |
| Non-Fat Milk (Blotting-Grade Blocker) | 1706404 | Bio-Rad |
| Nuclear Extraction kit | ab221978 | Abcam |
| Oligo-dT primers, 100 ul (0.4 ug/ul) | 79237 | Qiagen |
| Protease Inhibitor Cocktail Tablets | P8340 | Sigma-Aldrich |
| QIAamp DNA Mini Kit | 51304 | Qiagen |
| QIAzol Lysis Reagent | 57506291 | Qiagen |
| RiboGuard RNase inhibitor | E0126-40D7 | LGC Biosearch Technologies, Middleton, WI, USA |
| Senscript RT Kit Reagents | 205213 | Qiagen |
| T-PER™ Tissue Protein Extraction Reagent | 78510 | Thermo Fisher Scientific |
| TaqMan Universal Master Mix II, no UNG | 4440040 | Applied Biosystems, Waltham, MA, USA |

**Table S2. Primers for Kdm5c (X chromosome) and kdm5d (Y chromosome) genes used in fetal sex determination.**

| sex | primer | forward | reverse |
|---|---|---|---|
| *X-chromosome* | Kdm5c (F1) | 3'- CACCCCGGATCAGCATGTTT -5' | |
| *Y-chromosome* | Kdm5d(F2) | 3'-CCTTAGTCGGTAGAGTGGTT-5' | |
| *XY chromosome* | Kdm5c + kdm5d | | 5'- CCGCTGCCAAATTCTTTGG -3' |

We used the gene sequences of *Kdm5c* (X chromosome, NC_005120.4) and *Kdm5d* (Y chromosome, NC_024475.1) to determine fetal sex in rats.

**Table S3. Primer sequences for quantitative real-time PCR analysis of gene expression in placenta.**

| Gene | Sequence (5'→3') | Amplicon Size (bp) |
|---|---|---|
| ***il4*** | ***Forward:*** *CGTGATGTACCTCCGTGCTT* | 108 |
| | ***Reverse:*** *ATTCACGGTGCAGCTTCTCA* | |
| ***il6*** | ***Forward:*** *TGATGGATGCTTCCAAACTG* | 229 |
| | ***Reverse:*** *GAGCTTGGAAGTTGGGGTA* | |
| ***il1β*** | ***Forward:*** *CACCTTCTTTTCCTTCATCTTTG* | 241 |
| | ***Reverse:*** *GTCGTTGCTTGTCTCTCCTTGTA* | |
| ***il10*** | ***Forward:*** *CTGGCTCAGCACTGCTATGT* | 86 |
| | ***Reverse:*** *GCAGTTATTGTCACCCCGGA* | |
| ***tnfα*** | ***Forward:*** *ACTGAACTTCGGGGTGATTG* | 153 |
| | ***Reverse:*** *GCTTGGTGGTTTGCTACGAC* | |
| ***mmp8*** | ***Forward:*** *CCAAGGAGTGTCCAAGCCAT* | 127 |
| | ***Reverse:*** *TGCTAGTGGGGTAACCTGGA* | |
| ***mcp1*** | ***Forward:*** *CTGTCTCAGCCAGATGCAGTT* | 80 |
| | ***Reverse:*** *AGCCGACTCATTGGGATCA* | |

| | | |
|---|---|---|
| ***infγ*** | ***Forward:*** *GAGGAACTGGCAAAAGGACG*<br>***Reverse:*** *CAGGTGCGATTCGATGACAC* | 133 |
| ***f4/80*** | ***Forward:*** *GCCATAGCCACCTTCCTGTT*<br>***Reverse:*** *ATAGCGCAAGCTGTCTGGTT* | 143 |
| ***sod1*** | ***Forward:*** *TTGGCCGTACTATGGTGGTC*<br>***Reverse:*** *GGGCAATCCCAATCACACCA* | 120 |
| ***sod2*** | ***Forward:*** *CGGGGGCCATATCAATCACA*<br>***Reverse:*** *GCCTCCAGCAACTCTCCTTT* | 84 |
| ***catalase*** | ***Forward:*** *CTGACTGACGCGATTGCCTA*<br>***Reverse:*** *ATGGTGTAGGATTGCGGAGC* | 97 |
| ***gapdh*** | ***Forward:*** *AGACAGCCGCATCTTCTTGT*<br>***Reverse:*** *TACGGCCAAATCCGTTCACA* | 228 |

Primer sequences were designed using Primer-BLAST and validated for amplification efficiency prior to experimental analysis.

**Table S4. Primary and secondary antibodies used for Western blotting.**

| Target | Cat# | Vendor/City, State, Country | MW (kDa) | Dilution (primary antibody) | Secondary antibody (dilution; company, cat #) | Blocking buffer | RRIDs |
|---|---|---|---|---|---|---|---|
| MyD88(D80F5) | 4283 | Cell Signaling/Danvers, MA, USA | 33 | 1:250 | 1:2000; Anti -Rabbit IgG, HRP - Linked Antibody #7074, Cell Signaling | 3% BSA | RRID: AB_3073954 |
| NFkB p65 (L8F6) | 6956 | Cell Signaling/Danvers, MA, USA | 65 | 1:1000 | 1:5000; Anti-Mouse IgG, HRP- Linked Antiboby # 7076, Cell Signaling | 3% BSA | RRID: AB_397691 |
| TLR9 | PA5-20203 | Invitrogen/Waltham, MA, USA | 80 | 1:500 | 1:2000; Anti -Rabbit IgG, HRP - Linked | 3% Milk | RRID: AB_3073954 |

| | | | | | |
|---|---|---|---|---|---|
| | | | | | Antibody #7074, Cell Signaling |

**Table S5. ANOVA and pos-hoc output for Figure 3A**

| Endpoint | Interaction (Treatment × Sex) | Main effect: Treatment | Main effect: Sex | Sidak post hoc (Female: Saline vs. mtDNA) | Sidak post hoc (Male: Saline vs. mtDNA) | Sidak post hoc (Male vs. Female in Saline) |
|---|---|---|---|---|---|---|
| *il6* | F(1, 27)=10.58; p=0.0031 | F(1, 27)=10.61; p=0.003 | F(1, 27)=10.58; p=0.0031 | p>0.99 | p=0.0003 | p>0.99 |
| *il1b* | F(1, 27)=7.86; p=0.009 | F(1, 27)=39.3; p<0.0001 | F(1, 27)=99.6; p<0.0001 | p=0.056 | p<0.0001 | p<0.0001 |
| *ifnγ* | F(1, 15)=13.21; p=0.002 | F(1, 15)=10.23; p=0.006 | F(1, 15)=16.73; p=0.001 | p=0.0003 | p=0.98 | p=0.0002 |

| | | | | | | |
|---|---|---|---|---|---|---|
| *il4* | F(1, 27)=4.73; p=0.04 | F(1, 27)=62.06; p<0.0001 | F(1, 27)=252.7; p<0.0001 | p<0.0001 | p=0.0015 | p<0.0001 |
| *mcp1* | F(1, 27)=7.19; p=0.01 | F(1, 27)=111.6; p<0.0001 | F(1, 27)=36.06; p<0.0001 | p<0.0001 | p<0.0001 | p=0.084 |
| *tnf*a | F(1, 26)=0.802; p=0.378 | F(1, 26)=259.2; p<0.0001 | F(1, 26)=398.4; p<0.0001 | p<0.0001 | p<0.0001 | p<0.0001 |
| *il10* | F(1, 26)=0.097; p=0.757 | F(1, 26)=7.254; p=0.012 | F(1, 26)=542.9; p<0.0001 | p=0.124 | p=0.281 | p<0.0001 |
| *f4/80* | F(1, 26)=1.409; p=0.2460 | F(1, 26)=105.2; p<0.0001 | F(1, 26)=149.4; p<0.0001 | p<0.0001 | p<0.0001 | p<0.0001 |

| | | | | | | |
|---|---|---|---|---|---|---|
| *mmp8* | F(1, 26)=0.134; p=0.718 | F(1, 26)=0.157; p=0.695 | F(1, 26)=0.758; p=0.392 | p>0.99 | p=0.934 | p=0.977 |

Two-way ANOVA (factor 1, Treatment: saline, mtDNA; factor 2, Sex: male, female) followed by Sidak's multiple comparisons test. Planned comparisons: female saline vs. female mtDNA; male saline vs. male mtDNA; female saline vs male saline.

**Table S6. Multiplex panel of cytokine and chemokine levels in the placentas from male and female fetuses 4 h after exposure to purified mitochondrial DNA (mtDNA)**

| | Female | | Male | |
|---|---|---|---|---|
| **pg/mL** | Saline | mtDNA | Saline | mtDNA |
| **IL-1β** | 411.7 ± 55.44 | 454.3 ± 79.65 | 475.2 ± 112.2 | 405.8 ± 91.84 |
| **TNF-α** | 125.7 ± 71.97 | 76.90 ± 27.14 | 102.0 ± 42.43 | 93.77 ± 30.00 |
| **IL-4*** | 11.81 ± 9.56 | 7.55 ± 2.37 | 11.08 ± 6.91 | 5.30± 4.47 |
| **IL-10** | 13.12 ± 3.95 | 12.73 ± 4.76 | 12.64 ± 4.29 | 9.99 ± 4.22 |
| **MCP-1** | 594.7 ± 358.4 | 568.0 ± 322.8 | 941.2 ± 654.5 | 337.3 ± 158.5 |
| **CXCL1** | 1594 ± 484.3 | 1436 ± 545.5 | 1560 ± 397.1 | 1462 ± 494.7 |
| **RANTES*** | 974.4 ± 299.1 | 611.2 ± 187.3 | 727.3 ± 276.2 | 653.2 ± 283.7 |
| **M-CSF*** | 332.0 ± 98.22 | 218.4 ± 68.58 | 269.0 ± 80.33 | 171.5 ± 68.49 |

Protein concentrations of interleukin- (IL-) 1 beta (IL-1β), tumor necrosis factor-alpha (TNF-α), IL-4, IL-10, and chemokines monocyte chemoattractant protein (MCP-1), C-X-X motif chemokine ligand 1 (CXCL-1), regulated upon activation, normal T-cell expressed and secreted (RANTES), and macrophage colony stimulating factor (M-CSF) in placentas from female and male fetuses from pregnancies treated with saline or purified mitochondrial DNA (mtDNA). Data were analyzed by two-way ANOVA followed by Sidak's multiple comparisons test. *$p < 0.05$, treatment effect. Mean ± SD, n=7-8 placentas per sex-treatment group.

**Table S7. ANOVA and pos-hoc output for Figure 6A-F**

| Time-point | Endpoint | Interaction (Treatment × Sex) | Main effect: Treatment | Main effect: Sex | Sidak post hoc (Female: Saline vs. mtDNA) | Sidak post hoc (Male: Saline vs. mtDNA) | Sidak post hoc (Male vs. Female in Saline) |
|---|---|---|---|---|---|---|---|
| **4 h** | *sod1* | F(1, 26)=11.97; p=0.0019 | F(1, 26)=11.42; p=0.0023 | F(1, 26)=19.7; p=0.0001 | p=0.871 | p<0.0001 | p>0.99 |
| | *sod2* | F(1, 27)=5.22; p=0.03 | F(1, 27)=2.732; p=0.11 | F(1, 27)=29.08; p<0.0001 | p<0.0001 | p=0.115 | p=0.962 |
| | *catalase* | F(1, 25)=9.055; p=0.0059 | F(1, 25)=1.619; p=0.2149 | F(1, 25)=0.997; p=0.3276 | p=0.0305 | p=0.4044 | p=0.014 |

| | | | | | | | |
|---|---|---|---|---|---|---|---|
| **24 h** | *sod1* | F(1, 17)=0.08651; p=0.7722 | F(1, 17)=0.1398; p=0.7131 | F(1, 17)=0.004177; p=0.9492 | p=0.9937 | P=0.9974 | p>0.9999 |
| | *sod2* | F(1, 17)=19.38; p=0.0004 | F(1, 17)=0.4497; p=0.5155 | F(1, 17)=81.75; p<0.0001 | p=0.0726 | P=0.0035 | p<0.0001 |
| | *catalase* | F(1, 17)=10.84; p=0.0043 | F(1, 17)=0.6121; p=0.4448 | F(1, 17)=1.652; p=0.216 | p=0.309 | p=0.018 | p=0.4618 |

Two-way ANOVA (factor 1, Treatment: saline, mtDNA; factor 2, Sex: male, female) followed by Sidak's multiple comparisons test. Planned comparisons: female saline vs. female mtDNA; male saline vs. male mtDNA; female saline vs male saline.

**Table S8. Testing significance of Treatment effect.** $\chi^2$ test to compare deviance between full model and model excluding Treatment variable

| Model | ResDf | ResDev | Df | Dev | p-value |
|---|---|---|---|---|---|
| 1 | 13 | 19.642 | | | |
| 2 | 12 | 15.511 | 1 | 4.1306 | 0.04212 * |

*ResDf: residual degrees of freedom; ResDev: residual deviance; Df: degrees of freedom. *p-value < 0.05*

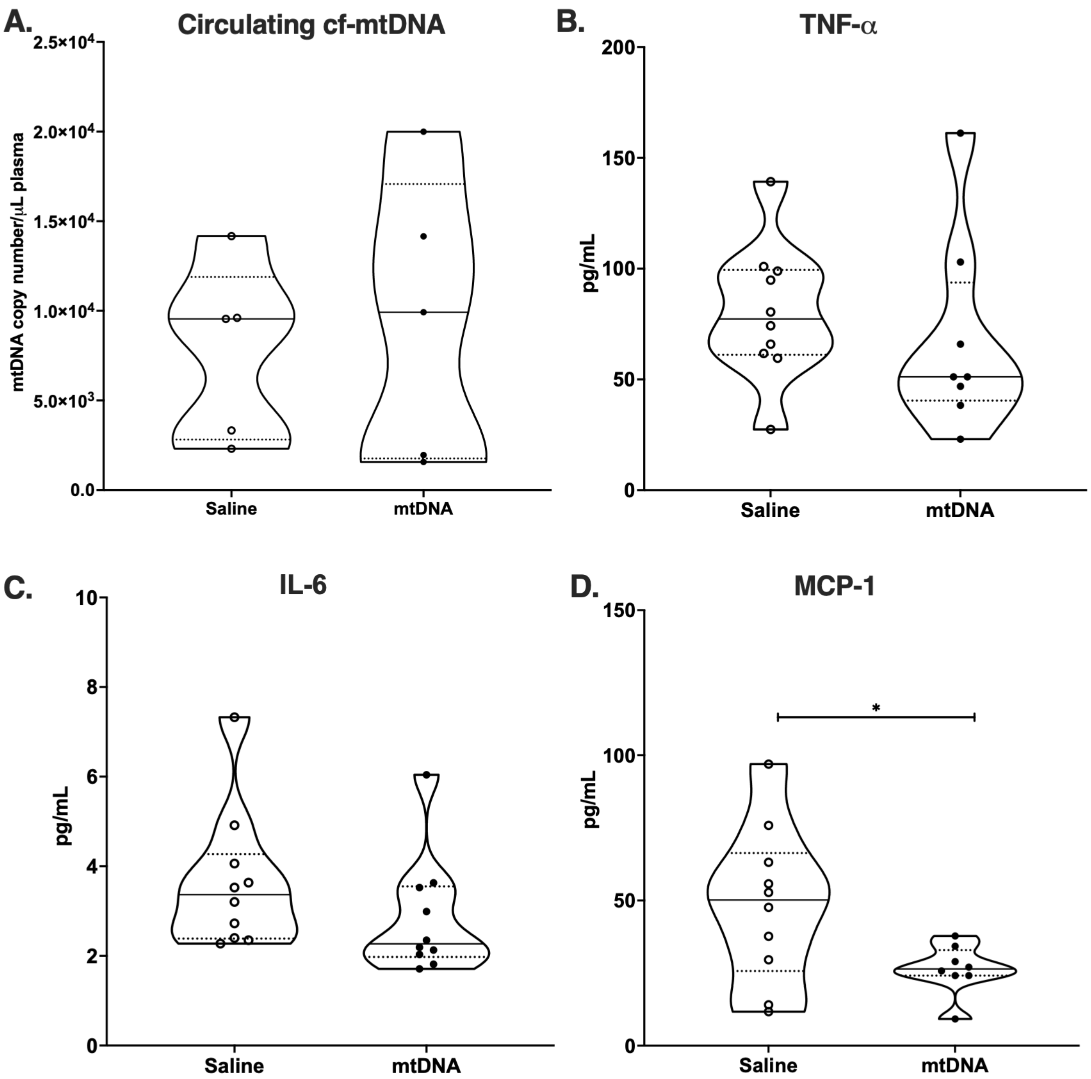

**Figure S1.** (A) Circulating cell-free mitochondrial DNA (mtDNA) copy number and (B) plasma concentrations of TNF-α, (C) IL-6, and (D) MCP-1 measured 4 h after saline or mitochondrial DNA (mtDNA) treatment in pregnant rats. Circulating mtDNA copy number did not differ between saline- and mtDNA-treated groups. Plasma TNF-α and IL-6 concentrations were not altered by mtDNA exposure, whereas MCP-1 concentrations were reduced in mtDNA-treated rats compared with saline-treated controls. Data are presented as means ± SD. Truncated violin plots show biological replicates, median (solid lines) and quartiles (dotted lines). Statistics were performed on log transformed mtDNA copy number data. mtDNA (n=5 dams per treatment group) and MCP-1 (n=10 dams per treatment group) data were analyzed using Welch's t-test, whereas IL-6 (n =10 dams per treatment group) and TNF-α (n = 8-10 dams per treatment group) data were analyzed using Mann-Whitney *U* test. *p<0.05.

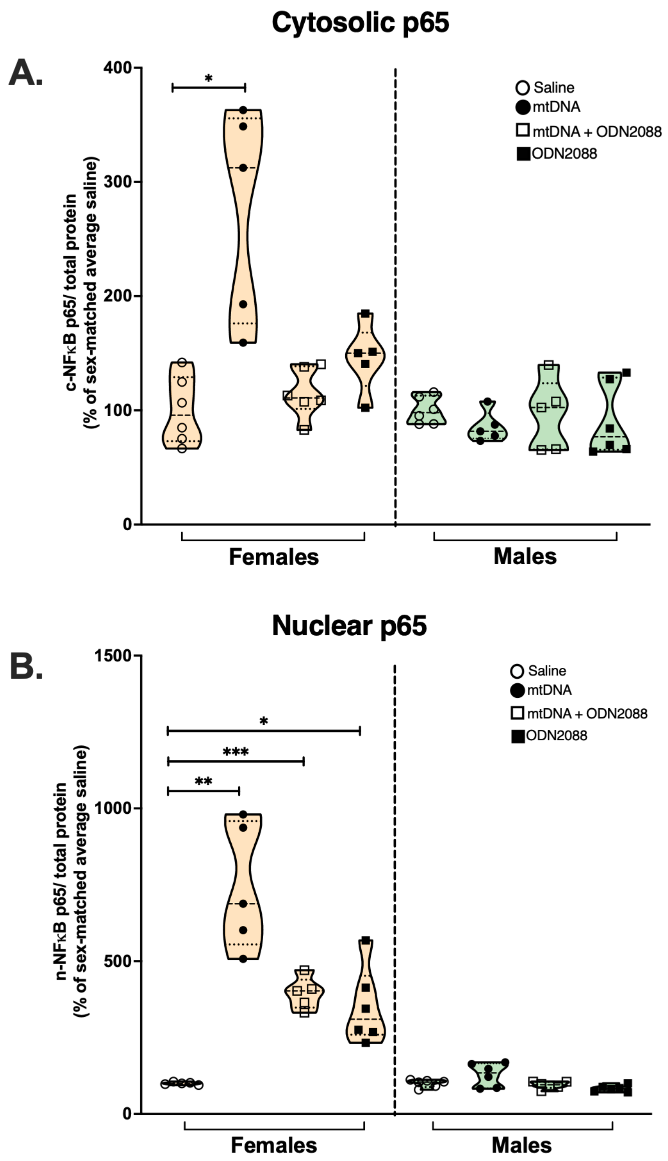

**Figure S2.** Nuclear factor kappa-light-chain-enhancer of activated B cells (NF-κB) expression in placentas from male and female fetuses after *in vivo* pharmacological antagonism of Toll-like receptor 9 (TLR9). Protein expression of NF-κB in (A) cytosolic and (B) nuclear fractions of placentas from female (orange) and male (green) fetuses of pregnant rats treated with saline (open circles), mitochondrial DNA (mtDNA; closed circles), mtDNA + TLR9 antagonist ODN2088 (open squares), or ODN2088 alone (closed squares). Protein levels were normalized to total protein and expressed relative to saline-treated controls (set to 100%). Welch ANOVA followed by Dunnett's T3 multiple comparisons was used for all datasets except cytosolic NF-κB in male placentas, which was analyzed with one-way ANOVA followed by Dunnett's multiple comparisons test. Truncated violin plots show individual placentas. One male and one female placenta were analyzed from each litter. Sample sizes are n = 5-6 placentas per sex-treatment group. Solid lines indicate medians and dotted lines indicate quartiles. *p<0.05, **p<0.01, ***p<0.001, ****p<0.0001.

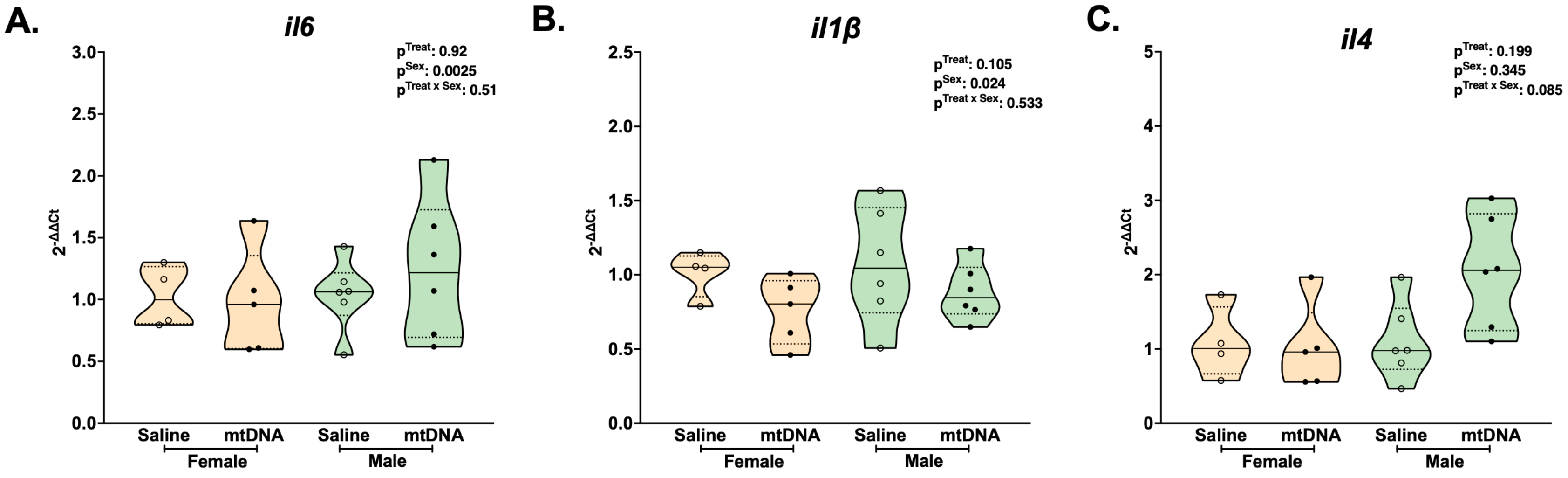

**Figure S3. Cytokine mRNA expression in rat placentas 24 h after exposure to purified mtDNA.** Relative mRNA expression ($2^{-\Delta\Delta Ct}$) of (A) interleukin-6 (*il6*), (B) *il1β*, and (C) *il4* in placentas from female (orange) and male (green) fetuses from pregnant rats treated with saline (open circles) or mitochondrial DNA (mtDNA, closed circles) extracted from rat liver. Statistics were performed on ΔCt values using two-way ANOVA followed by Sidak's multiple comparisons test. Truncated violin plots show individual placentas. One male and one female placenta were analyzed from each litter. Sample sizes are n = 4-6 placentas per sex-treatment group. Solid lines indicate medians and dotted lines indicate quartiles.

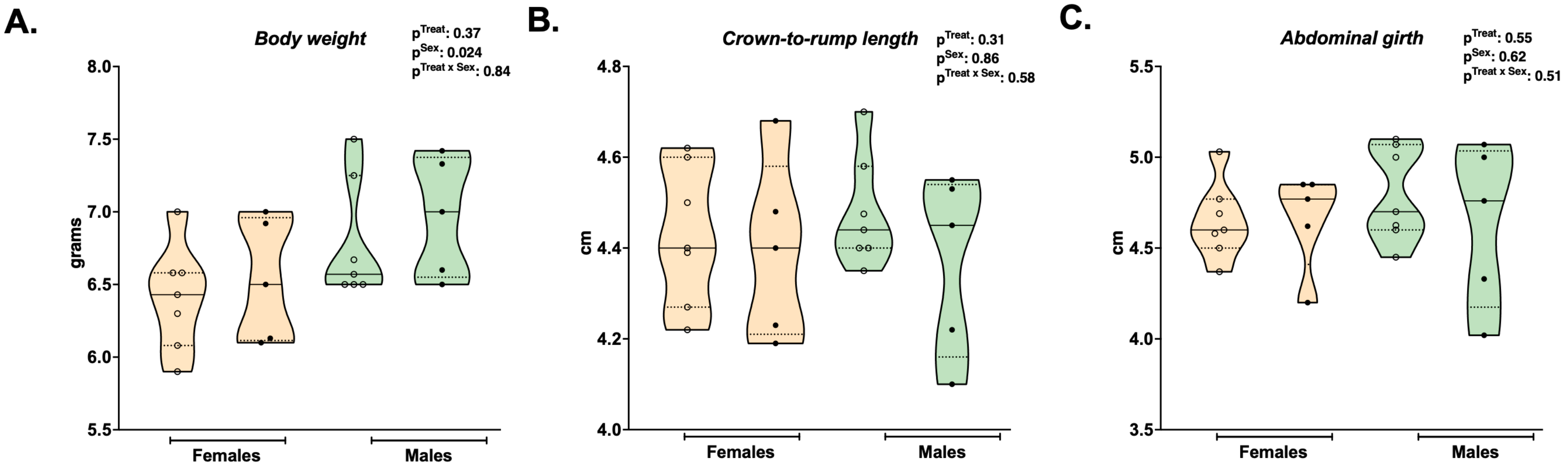

**Figure S4. Biometrics of neonates from pregnancies exposed to an acute mtDNA challenge during pregnancy.** (A) Body weight, (B) crown-to-rump length, and (C) abdominal girth of female (orange) and male (green) neonates from pregnancies exposed to saline (open circles) or mitochondrial DNA (mtDNA, closed circles). Statistics were performed using two-way ANOVA followed by Sidak's multiple comparisons test. Truncated violin plots show individual placentas. One male and one female placenta were analyzed from each litter. Sample sizes are n = 5-7 placentas per sex-treatment group. Solid lines indicate medians and dotted lines indicate quartiles.
