## Supplementary material for "Fetal sex shapes placental inflammatory responses to extracellular mitochondrial DNA": Blot images

**SUPPLEMENTARY FILE 2: PCR GEL IMAGES FOR SEX DETERMINATION AND WESTERN BLOT IMAGES**

### Fetal sex shapes placental inflammatory responses to extracellular mitochondrial DNA

Reneé de Nazaré Oliveira da Silva[1], Nataliia Hula[1], Desirae Escalera[1], Leslie Lopez [1], Gabrielle Kelly[1], Isabelle K. Gorham[2], Megan Rowe[2], Contessa A. Ricci[3], Ciprian Gheorghe[1,4], Nicole R. Phillips[2], Styliani Goulopoulou[1, 4]

[1]Lawrence D. Longo, MD Center for Perinatal Biology, Department of Basic Sciences, Loma Linda University, Loma Linda, California, USA; [2]Department of Microbiology, Immunology, and Genetics, University of North Texas Health Science Center, Fort Worth, Texas, USA; [3]Institute for Research and Education to Advance Community Health (IREACH), College of Nursing, Elson S. Floyd College of Medicine, Washington State University, USA; [4]Department of Gynecology and Obstetrics, Loma Linda University, Loma Linda, California, USA

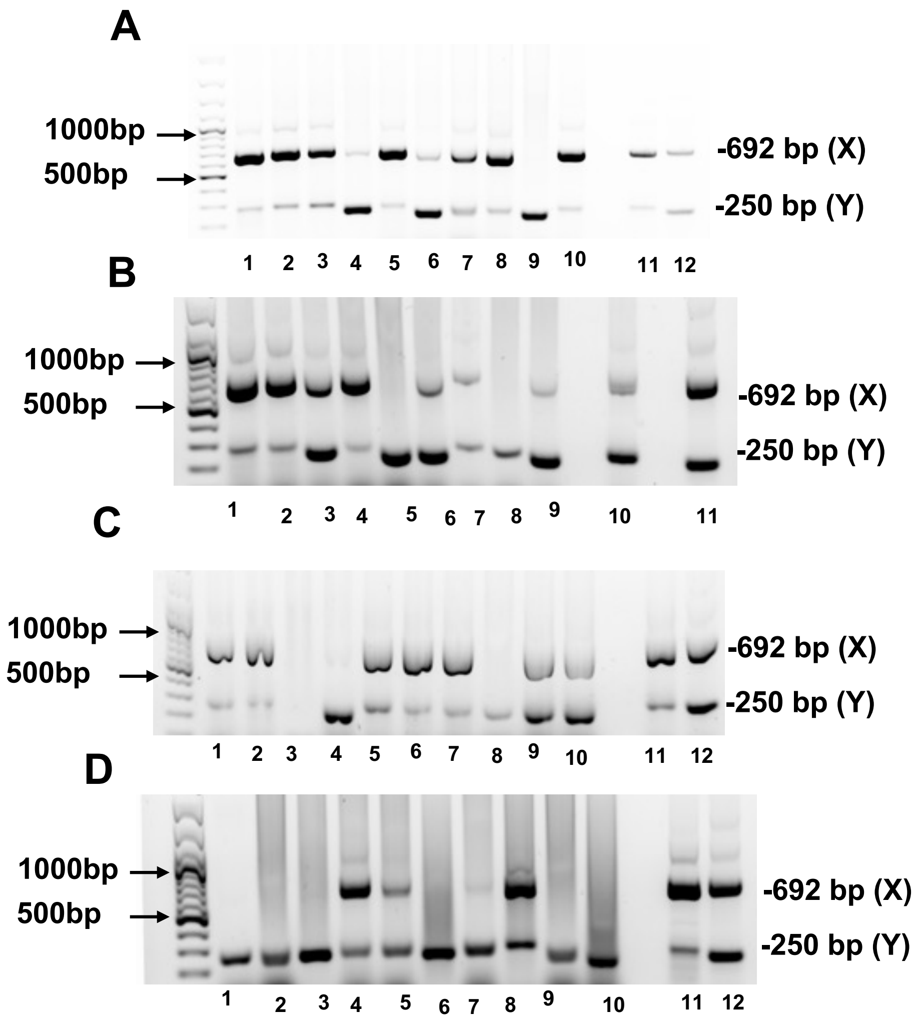


**Fetal PCR products corresponding to X and Y chromosome-specific genes (Part 1)**. Each panel represents a different gel loaded by samples from different animals. We identified males by a band at 250 and 692 bp, and females by a band at 692 bp. Bands were viewed on 1% agarose gel using GeneRuler 100-bp Plus DNA Ladder. Tails from adult male and female rats were used as controls. **(A) Saline:** Lanes 1-10: fetal samples from saline-treated pregnancies, Lane 11: Female adult rat tail, control; Lane 12: Male adult rat tail, control; **(B) Saline:** Lane 1: Female adult rat tail, control; Lanes 2-10: fetal samples from saline-treated pregnancies, Lane 11: Male adult rat tail, control; **(C) mtDNA:** Lanes 1-10: fetal samples from mtDNA-treated pregnancies, Lane 11:

Female adult rat tail, control, Lane 12: Male adult rat tail, control; **(D) mtDNA:** Lanes 1-10: fetal samples from mtDNA-treated pregnancies, Lane 11: Female adult rat tail, control, Lane 12: Male adult rat tail, control.

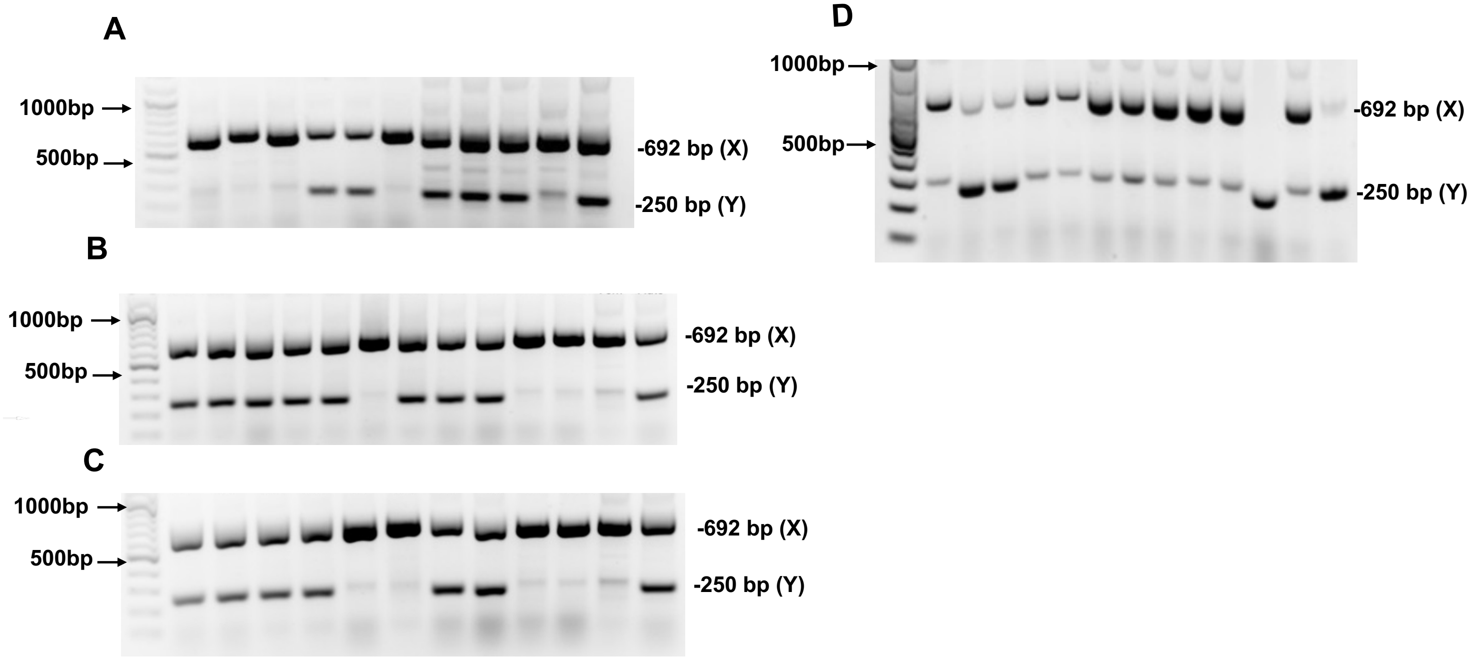


**Fetal PCR products corresponding to X and Y chromosome-specific genes (Part 2)**. Each panel represents a different gel loaded by samples from different animals. We identified males by a band at 250 and 692 bp, and females by a band at 692 bp. Bands were viewed on 1% agarose gel using GeneRuler 100-bp Plus DNA Ladder. Tails from adult male and female rats were used as controls. **(A) mtDNA+ODN2088:** Lanes 1-9: fetal samples from mtDNA+ODN2088-treated pregnancies, Lane 10: Female adult rat tail, control, Lane 11: Male adult rat tail, control; **(B) ODN2088:** Lanes 1-11: fetal samples from ODN2088-treated pregnancies, Lane 12: Female adult rat tail, control, Lane 13: Male adult rat tail, control; **(C) mtDNA+ODN2088:** Lanes 1-10: fetal samples from mtDNA+ODN2088-treated pregnancies ,Lane 11: Female adult rat tail, control; Lane 12: Male adult rat tail, control; **(D) ODN2088:** Lanes 1-11: fetal samples from ODN2088-treated pregnancies, Lane 12: Female adult rat tail, control; Lane 13: Male adult rat tail, control.

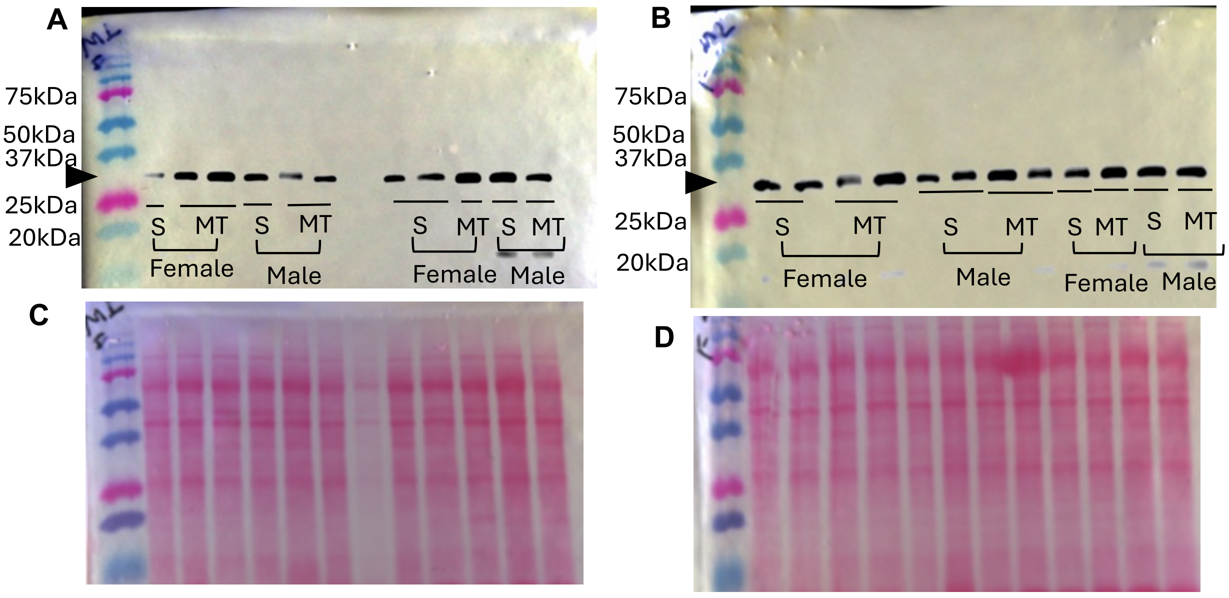


Representative immunoblot staining against (A, B) myeloid differentiation primary response 88 (MyD88) and (C, D) Ponceau S total protein staining in cytosolic placental fractions from saline-treated (S) and mitochondrial DNA-treated (MT) rats. Placental samples correspond to placentas from female and male fetuses.

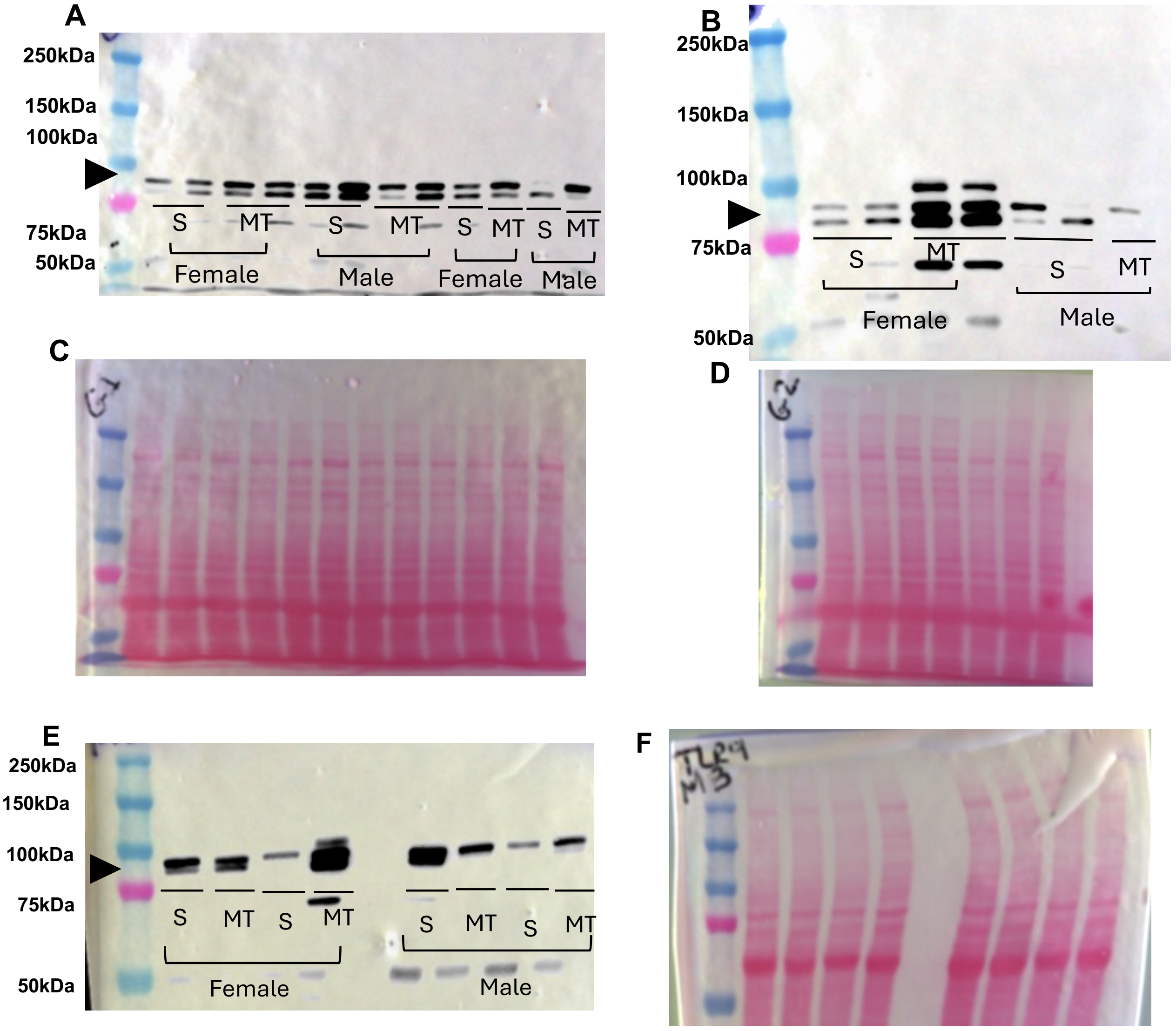


Representative immunoblot staining against (A, B, E) Toll-like receptor 9 (TLR9) and (C, D, F) Ponceau S total protein staining in cytosolic placental fractions from saline-treated (S) and mitochondrial DNA-treated (MT) rats. Placental samples correspond to placentas from female and male fetuses.

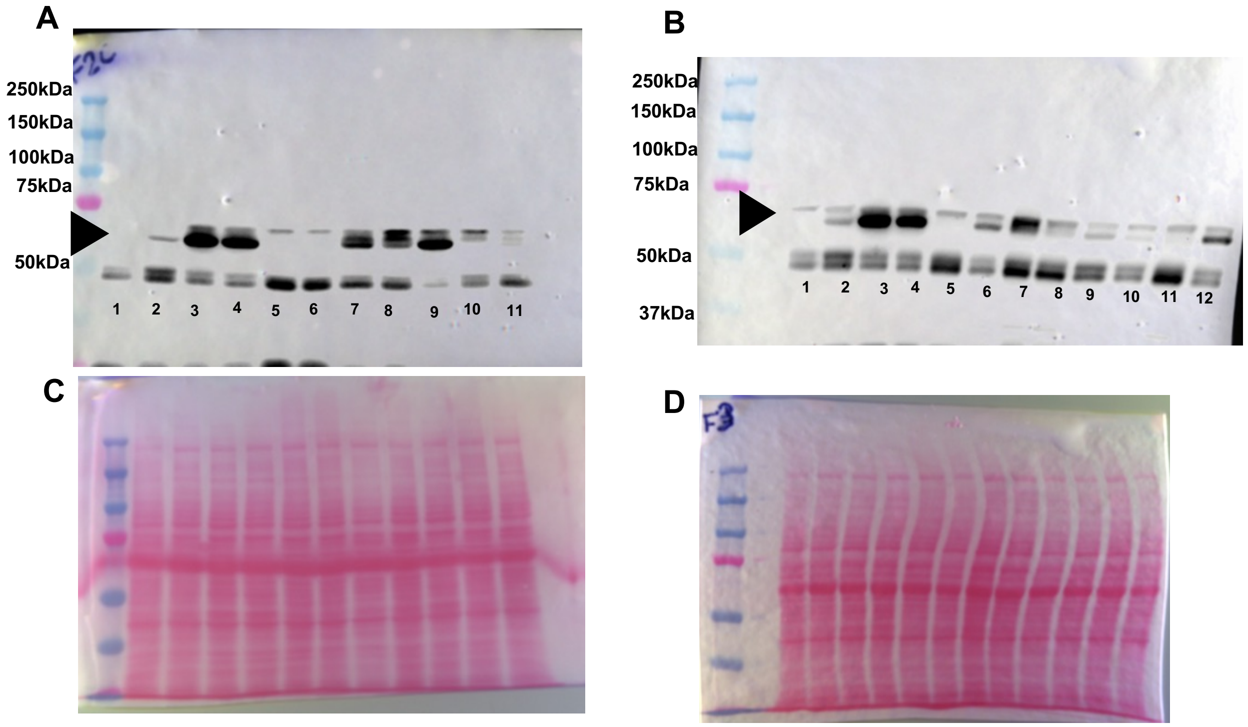


Representative immunoblot staining against (A, B) Nuclear factor kappa-light-chain-enhancer of activated B cells (NF-κB) and (C, D) Ponceau S total protein staining in cytosolic placental fractions from saline-treated (1, 2 ,8), mitochondrial DNA-treated (3, 4, 9), mitochondrial DNA plus ODN2088-treated (5, 6, 10), and ODN2088 plus saline-treated (7, 11, 12) rats. Placental samples correspond to placentas from female fetuses.

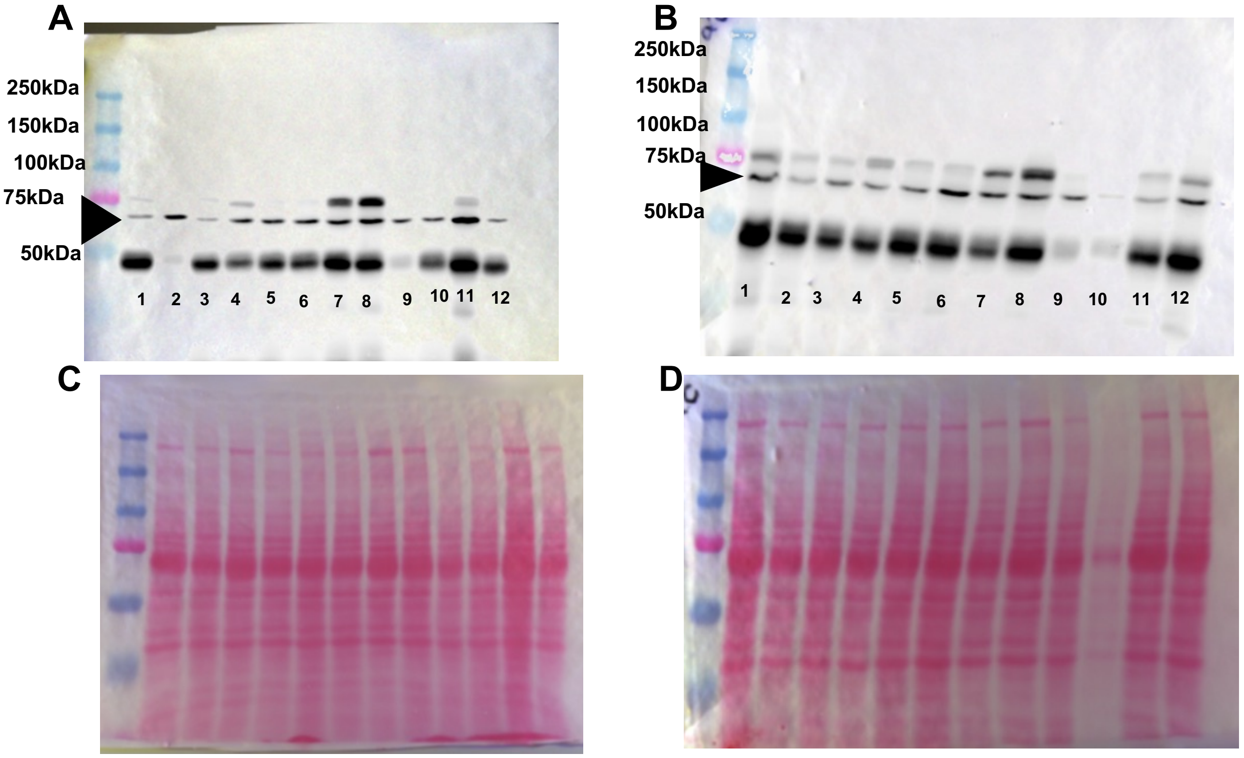


Representative immunoblot staining against (A, B) Nuclear factor kappa-light-chain-enhancer of activated B cells (NF-κB) and (C, D) Ponceau S total protein staining in cytosolic placental fractions from saline-treated (1, 2, 9), mitochondrial DNA-treated (3, 4, 10), mitochondrial DNA plus ODN2088-treated (5, 6, 11), and ODN2088 plus saline-treated (7, 8, 12) rats. Placental samples correspond to placentas from male fetuses.

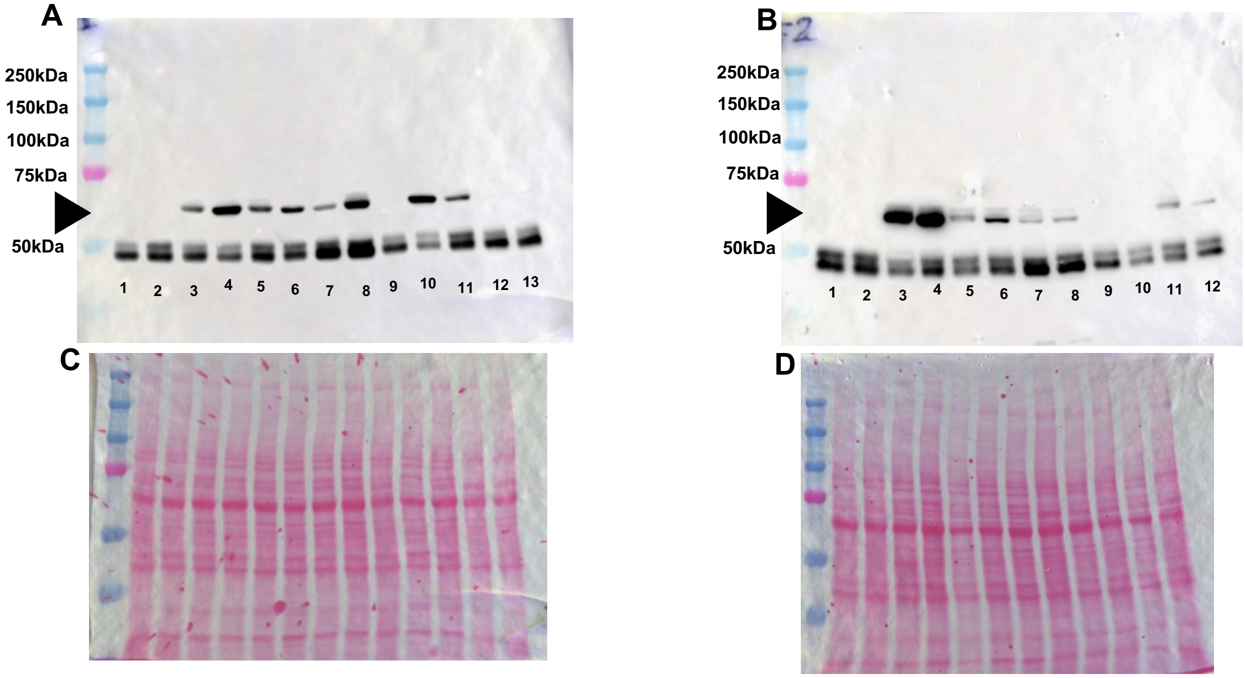


Representative immunoblot staining against (A, B) Nuclear factor kappa-light-chain-enhancer of activated B cells (NF-κB) and (C, D) Ponceau S total protein staining in nuclear placental fractions from saline-treated (1, 2, 9), mitochondrial DNA-treated (3, 4, 10), mitochondrial DNA plus ODN2088-treated (5, 6, 11), and ODN2088 plus saline-treated (7, 8, 12, 13) rats. Placental samples correspond to placentas from female fetuses.

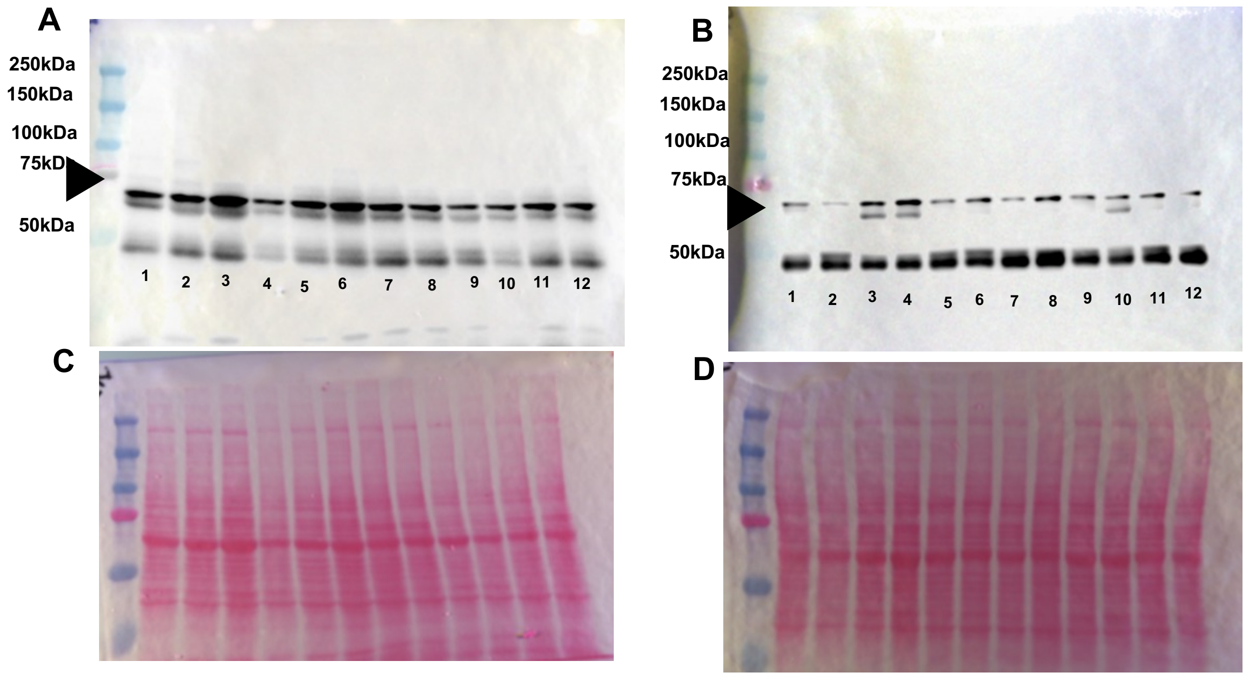


Representative immunoblot staining against (A, B) Nuclear factor kappa-light-chain-enhancer of activated B cells (NF-κB) and (C, D) Ponceau S total protein staining in nuclear placental fractions from saline-treated (1, 2, 9), mitochondrial DNA-treated (3, 4, 10), mitochondrial DNA plus ODN2088-treated (5, 6, 11), and ODN2088 plus saline-treated (7, 8, 12) rats. Placental samples correspond to placentas from male fetuses.
